# Identification of a fungal antibacterial endopeptidase that modulates immune responses

**DOI:** 10.1101/2024.09.13.612890

**Authors:** Silke Machata, Ute Bertsche, Franziska Hoffmann, Zaher M. Fattal, Franziska Kage, Michal Flak, Alexander N. J. Iliou, Falk Hillmann, Ferdinand von Eggeling, Hortense Slevogt, Axel A. Brakhage, Ilse D. Jacobsen

## Abstract

*Aspergillus fumigatus* is a saprophytic fungus dwelling in soil and on decaying plant material, but also an opportunistic pathogen in immunocompromised patients. In its environmental niche, *A. fumigatus* faces competition from other microorganisms including bacteria. Here, we describe the discovery of the first secreted antibacterial protein in *A. fumigatus*. We identified a secreted fungal endopeptidase, designated CwhA, that cleaves peptidoglycan of Gram-positive bacteria at specific residues within the peptidoglycan stem peptide. Cleavage leads to bacterial lysis and the release of peptidoglycan cleavage products. Expression of *cwhA* is induced by the presence of bacteria. Furthermore, CwhA is highly abundant in murine lungs during invasive pulmonary aspergillosis and peptidoglycan cleavage products generated by CwhA stimulate cytokine production of human immune cells. Although CwhA does not affect human cells directly, this novel player in fungal-bacterial interactions could affect *A. fumigatus* infections by inhibiting Gram-positive bacteria in its vicinity, and modulating the immune system.

## Introduction

Saprophytic filamentous fungi such as *Aspergillus* (*A.*) *fumigatus* share their environmental niche in soil and on decaying plant material with a plethora of other microorganisms. In order to survive in this highly competitive environment, filamentous fungi produce a variety of secondary metabolites, including antibiotics, whose production is often induced by the presence of other microorganisms (Adnani *et al*, 2017; Fischer *et al*, 2018; Netzker *et al*, 2015; Netzker *et al*, 2018; Schroeckh *et al*, 2009). Some of these molecules also play a role in infections with *A. fumigatus* that occur in immunocompromised patients and patients with underlying lung disease such as cystic fibrosis (CF). One example is gliotoxin, a mycotoxin with amoebicidal properties that negatively affects immune cells and thereby contributes to virulence (Hillmann *et al*, 2015a; Kosmidis & Denning, 2015; Scharf *et al*, 2016). Another example is the *A. fumigatus* pigment DHN-melanin that affects phagocytosis and killing of the fungus by both environmental amoeba and mammalian macrophages (Akoumianaki *et al*, 2016; Ferling *et al*, 2020).

Polymicrobial communities are not only found in the environment but also in human and animal hosts. Probably best studied is the complex network of the gastrointestinal microbiota and mycobiota, which was shown to have a strong impact on human health (Krüger *et al*, 2019). In contrast, microorganisms colonizing the lower respiratory tract have been largely neglected in the past and the healthy lung was long considered to be a sterile environment (Barcik *et al*, 2020; Levy *et al*, 2017; Man *et al*, 2017). This view has changed due to the advances in culture-independent techniques for microbiological analysis, recognizing the complexity of respiratory microbial communities (Chang *et al*, 2020). Characterized by comparatively low microbial density (10^3^/g of lung tissue) the lung microbiome composition was shown to change drastically in case of chronic lung diseases such as CF or asthma, leading to dysbiosis and pulmonary infections (Chotirmall & McElvaney, 2014; Dickson *et al*, 2015; Mathieu *et al*, 2018; Mitchell & Glanville, 2018). In CF patients, polymicrobial infections often occur that vary in composition and diversity depending on the patient’s age. The majority of adult CF patients’ lungs are colonized by *Staphylococcus* (*S.*) *aureus*, the *Burkholderia cepacia* complex, and *Pseudomonas* (*P.*) *aeruginosa* (Blanchard & Waters, 2019). Among fungi, *A. fumigatus* and *Candida albicans* are the most prevalent colonizers in the airways of CF patients (Chotirmall & McElvaney, 2014). In this setting, interactions between *A. fumigatus* and colonizing bacteria are likely (Poore *et al*, 2021) and have been studied in some detail for *P. aeruginosa*. *P. aeruginosa* can suppress growth of *A. fumigatus* by various mechanisms including phenazines, secondary metabolites that also act as virulence factors, and siderophore-mediated iron depletion (Briard *et al*, 2015; Keown *et al*, 2020). In addition, *Klebsiella pneumoniae* was reported to inhibit spore germination and hyphal development of *Aspergillus* spp. *in vitro* (Nogueira *et al*, 2019). The defence mechanisms of *Aspergillus* against bacterial competitors are less well studied. It was shown that the *A. fumigatus* secondary metabolite gliotoxin inhibits *P. aeruginosa* in co-culture experiments (Reece *et al*, 2018). Moreover, a recent publication describes the discovery of afusins, defensin-like peptides in *A. fumigatus* which are induced during conidiation and mediate protection of conidia against both Gram-positive and Gram-negative bacteria (Dümig *et al*, 2021). However, no reports of proteins produced by *A. fumigatus* mycelia that could play a role in multispecies interaction by suppressing bacterial growth exist thus far.

In our study presented here, we describe an antibacterial fungal protein, designated CwhA, which is highly abundant in lungs of mice infected with *A. fumigatus* (Machata *et al*, 2020). We demonstrate that *cwhA* gene expression is induced by the presence of bacteria and during late stages of infection of alveolar epithelial cells. The protein acts as an endopeptidase on peptidoglycan (PG) of Gram-positive bacteria, leading to bacterial lysis and the release of a PG cleavage product that stimulates cytokine production of human immune cells. This novel player in fungal-bacterial interactions could affect *A. fumigatus* infections by inhibiting Gram-positive bacteria in its vicinity, and modulating the immune system.

## Results

### The *A. fumigatus* protein CwhA is produced during infection and contains functional domains of bacterial cell wall hydrolases

Several fungal proteins of *A. fumigatus* were recently identified in bronchoalveloar lavage (BAL) samples of mice with invasive aspergillosis (Machata *et al*., 2020). One of the most highly abundant proteins was the uncharacterized fungal protein B0YAY0 (AFUB_086210). This protein has not been reported as a secreted protein in previous proteome studies of *A. fumigatus* grown *in vitro* but the respective gene was shown to be induced in a mouse infection model by transcriptome analysis (McDonagh *et al*, 2008; Schwienbacher *et al*, 2005; Wartenberg *et al*, 2011). We confirmed the production of B0YAY0 during invasive pulmonary aspergillosis (IPA) in mice by MALDI-imaging mass spectrometry (MSI) of lung slices: Six m/z values were detected by MSI which correlated to B0YAY0 peptide fragments predicted by *in silico* digestion (Supplementary Table 1). These m/z - values were also detected in control spots containing the recombinant protein. As an example, the distribution of one peptide with the mass m/z 1174.334 corresponding to the peptide (K)KEYNTLECR(G) [M+NH4]^+^ is shown in Figure 1. The protein was heterogeneously distributed in areas of hyphal growth, implying secretion of the protein from the fungal mycelia (Figure 1). According to *in silico* analysis the protein contains a signal peptide responsible for protein secretion (1-35 aa), a bacterial SH3 domain (Pfam PF08239, 21-97 aa), and a protein domain of the NlpC/p60 family (Pfam PF00877, 141-247 aa) (Figure 2A). Proteins of this superfamily have been mainly described in bacteria and exhibit functional diversity (Anantharaman & Aravind, 2003). The NlpC/p60 domain carries a catalytic triad that consists of a cysteine, a histidine and a polar residue and is often found in cell wall hydrolases of bacteria (Machata *et al*, 2005; Vermassen *et al*, 2019; Xu *et al*, 2015). NlpC/p60 cell wall hydrolases can act as lytic enzymes that permeabilize the bacterial peptidoglycan (PG) layer and potentially lead to lysis. Due to its structural similarity with bacterial cell wall hydrolases, the protein B0YAY0 was designated CwhA (cell wall hydrolase A) in this study.

**Figure 1.**
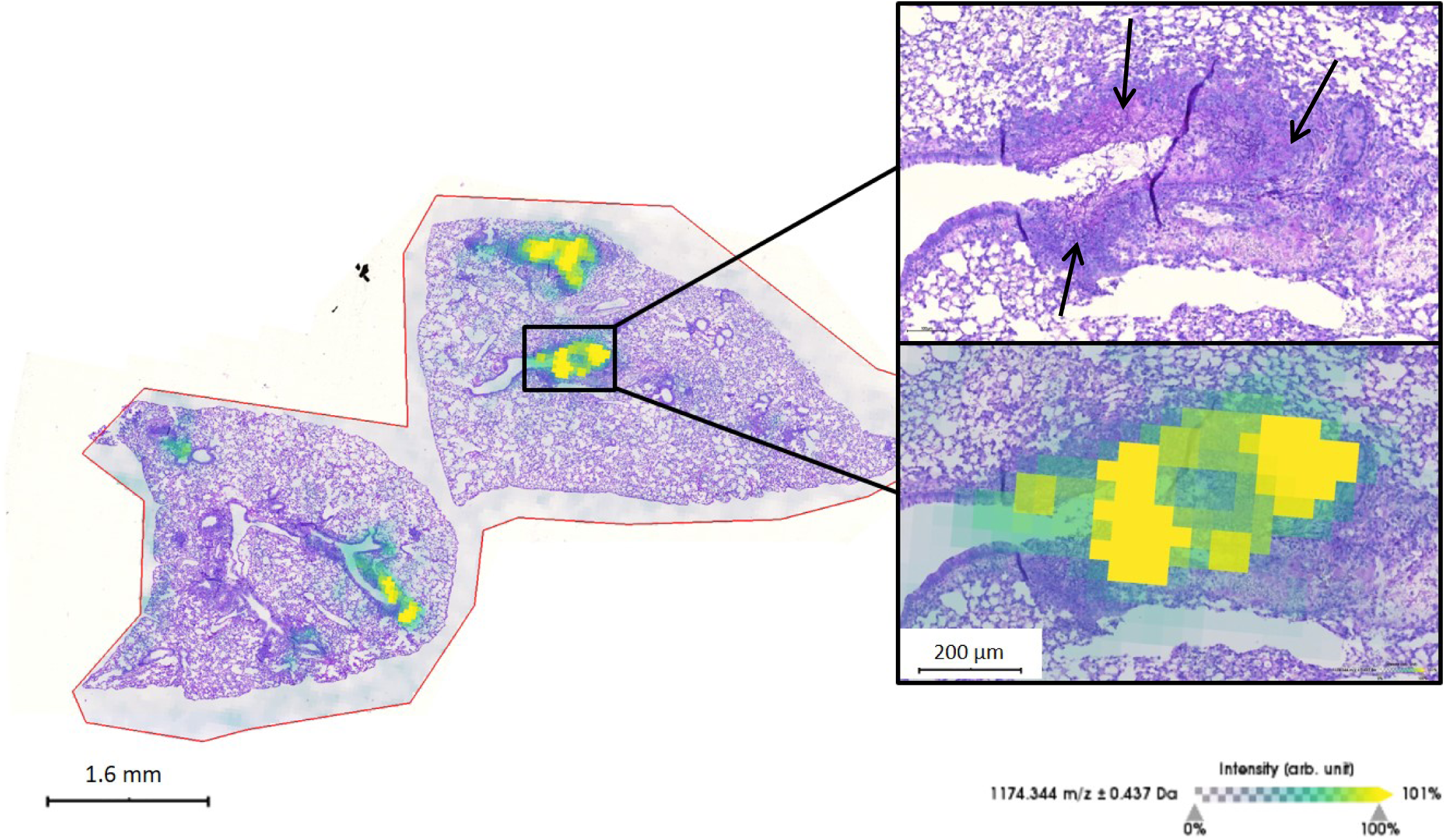
Detection of the secreted *A. fumigatus* protein B0YAY0 (CwhA) in infected murine lungs by MALDI-imaging. Representative lung histolology of leukopenic mice infected with *A. fumigatus.* Fungal mycelia are visualized with periodic acid Schiff staining and marked by black arrows in the upper enlarged image. The fungal protein was detected as a mass of m/z 1174.704 corresponding to the peptide (K)KEYNTLECR(G) with a mass of 1156.311 [M+NH4]. The lower enlarged image displays relative abundance of the fungal protein as colored areas.

**Figure 2.**
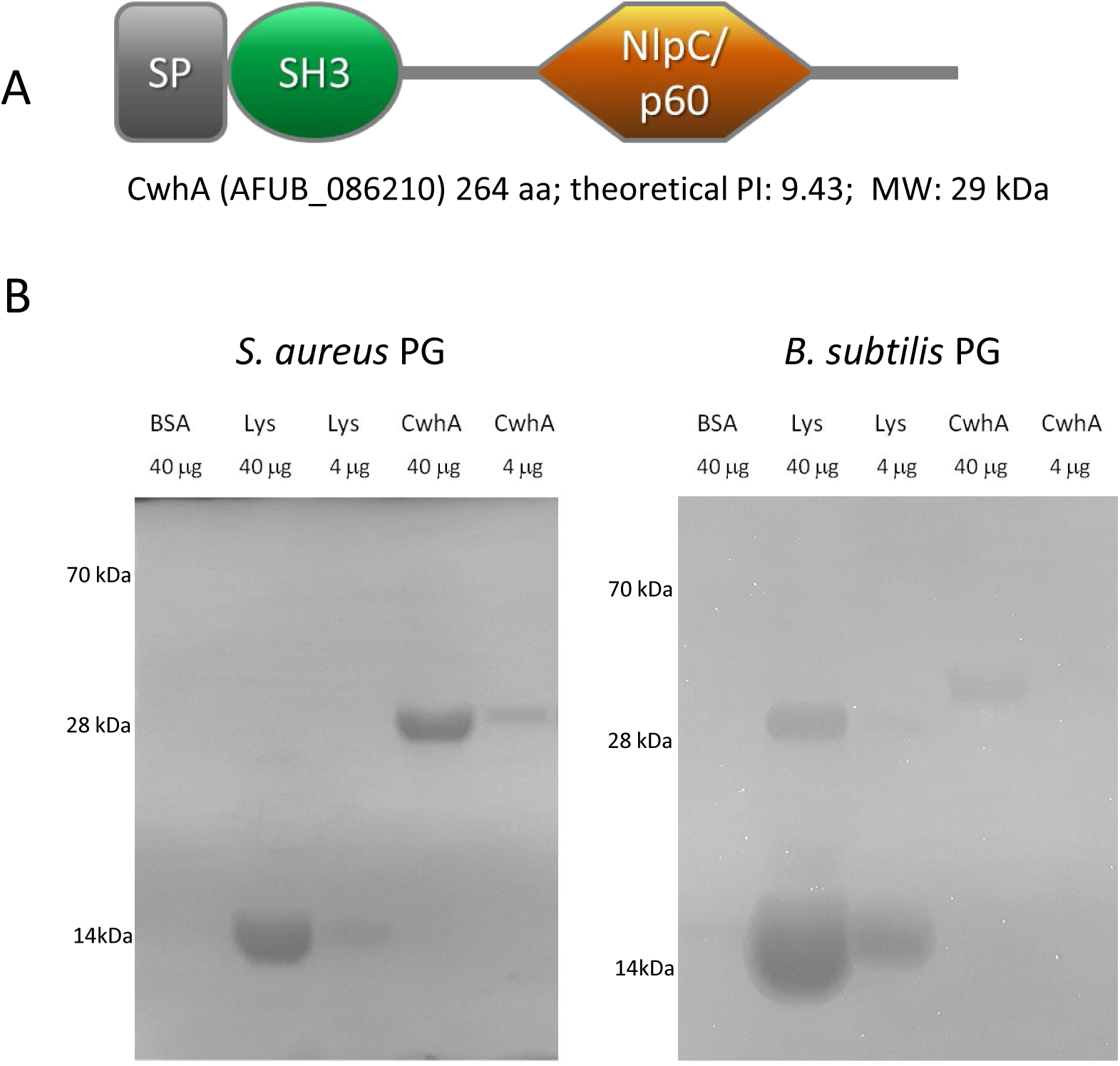
Predicted protein domains and visualization of cell wall hydrolytic activity of ChwA. (A) Protein domains as predicted by Pfam; *A. fumigatus* CwhA contains a signal peptide (SP), SH3 bacterial domain (PF08239) and NlpC/p60 family domain (PF00877). (B) Hydrolytic activity of CwhA against peptidoglycan (PG) from *S. aureus* and *B. subtilis* was visualized by zymography. CwhA, BSA (negative control) and lysozyme (Lys; positive control) were applied in the indicated concentrations to zymogram gels containing bacterial cell wall substrate. Lytic activity leads to clear bands in blue-stained gels. The pictures are inverted to mark cleavage products as dark bands. Representative images of three to four independent experiments using two independent protein preparations are shown.

### *A. fumigatus* CwhA cleaves bacterial PG

To determine if CwhA has hydrolytic activity against bacterial PG, we produced a recombinant protein by replacing the signal peptide sequence with an N-terminal His-tag. This protein fusion was produced in *Escherichia coli* and purified by affinity chromatography. The recombinant protein was tested for its activity using zymogram gels containing PG from *S. aureus* and *Bacillus* (*B.*) *subtilis* as substrates (Figure 2B). Cleavage of PG by the activity of the protein resulted in a band with an expected molecular mass of 29 kDa. CwhA was highly active against cell wall material from *S. aureus* and moderately active against *B. subtilis* (Figure 2B). In comparison, the control enzyme lysozyme showed strong activity against *B. subtilis* and was less active against *S. aureus* PG. Based on these findings we conclude that CwhA acts as a cell wall hydrolase against bacterial PG with particular substrate preference.

### CwhA promotes lysis of Gram-positive bacteria

To assess the impact of the protein on live bacteria, we treated bacterial cultures of *S. aureus* with recombinant CwhA at various concentrations (16 to 200 µg/ml) and tested bacterial growth by measuring the optical density (OD). While bacterial growth was unaffected when CwhA was added to LB media (Supplementary Figure 1), we observed a clear decrease of the OD when bacteria were suspended in Tris buffer (50 mM Tris pH 8.0) and treated with CwhA (Figure 3A). Due to the hypoosmotic conditions in the buffer, damaging effects on the bacterial cell wall are likely to have a stronger impact on the integrity of the bacterial cell compared to complex growth media. Already 10 minutes upon exposure to CwhA, the OD was considerably reduced compared to the PBS control, showing progressively stronger effects with increasing protein concentrations. Within two hours of treatment, the OD dropped between 30 % and 50 % in a dose-dependent manner, while a decrease of only 10 % could be observed in untreated bacterial cells in the PBS control (Figure 3A). In addition to the *S. aureus* strain SA113, we tested methicillin-resistant *S. aureus* strains (MRSA) for their susceptibility to CwhA. CwhA reduced the OD of these MRSA strains to a similar extend as observed for the SA113 reference strain (Figure 3B and Supplementary table 2). Other Gram-positive bacteria, such as *Enterococcus faecalis* and *Streptococcus pneumoniae,* were also susceptible to CwhA-mediated lysis, while the Gram-negative bacteria *Pseudomonas aeruginosa* and *Klebsiella pneumoniae* were resistant (Supplementary Figure 2). In contrast to the other tested Gram-positive bacteria, we observed no OD decrease of *B. subtilis* upon treatment of the culture with CwhA (Supplementary Figure 2). This observation coincides with the zymogram data showing only weak activity of CwhA against *B. subtilis* cell wall substrate and suggests a substrate preference of CwhA for murein from *S. aureus* over *B. subtilis* or Gram-negative bacteria.

**Figure 3.**
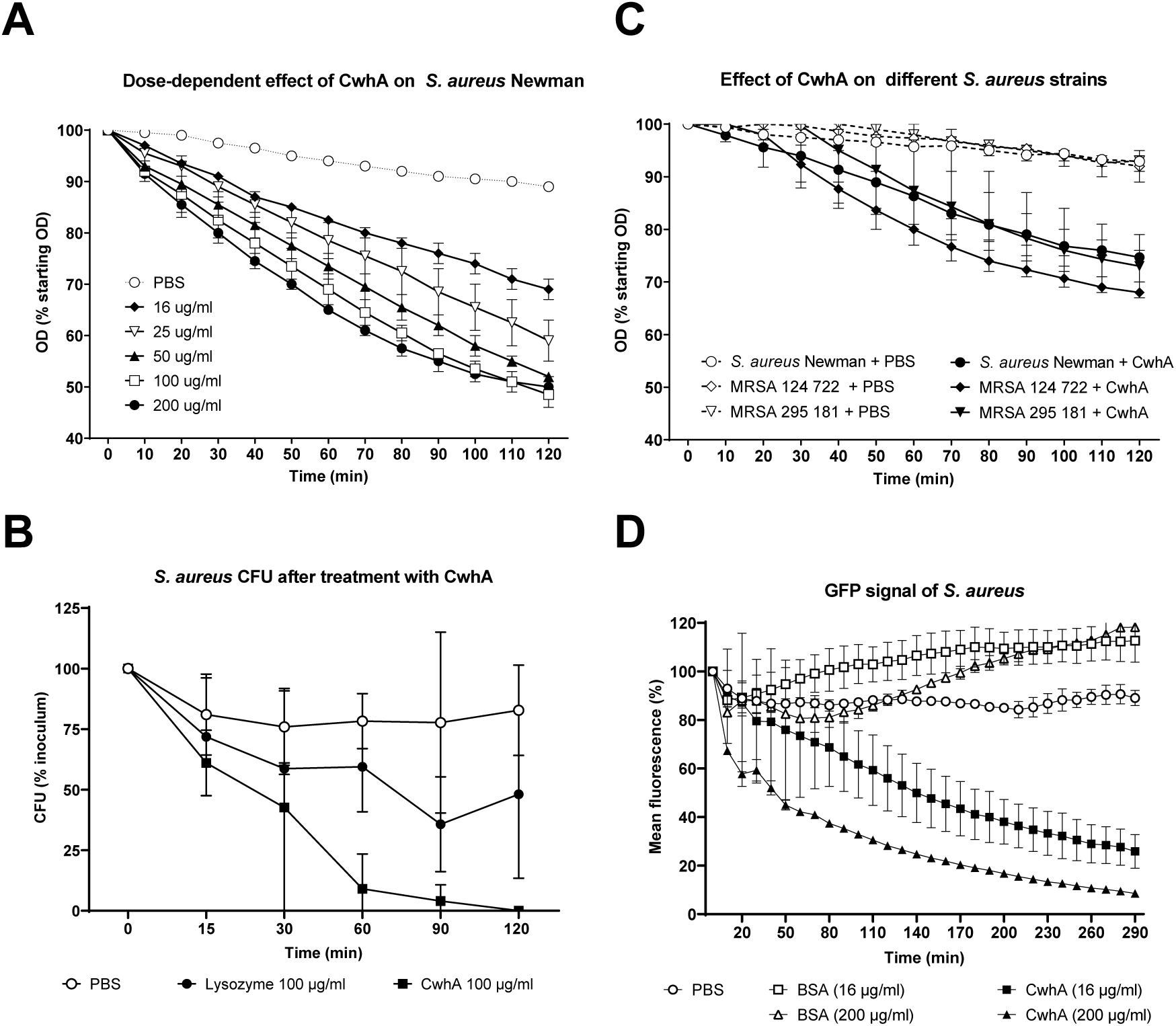
Effect of recombinant CwhA on *S. aureus*. (A) The optical density (600 nm) after addition of recombinant CwhA (16-200 μg/ml) to bacterial suspensions in 50 mM Tris buffer (pH 8) was measured over time; two independent experiments, data shown as mean and range. (B) Decrease of optical density of *S. aureus* strain Newman and two MRSA strains after addition of 100 μg/ml CwhA (three independent experiments, data shown as mean ± SD). Data was statistically analyzed by 2-way ANOVA comparing the control with treatment for each strain. A significant treatment effect was observed for strains Newman (p=0.0373) and 124 722 (p=0.0022). (C) The colony forming units (CFU) of *S. aureus* were determined at the indicated time points after addition of PBS, 100 μg/ml CwhA or 100 μg/ml lysozyme. Data from three independent experiments (except 120 min time point: two experiments) are presented as mean ± SD. (D) GFP-signal of a GFP-expressing *S. aureus* strain after treatment with CwhA. Bovine serum albumin (BSA) and PBS were used as controls. Two independent experiments, data shown as mean and range. Time elapse videos of *S. aureus*::Gfp with and without CwhA treatment are supplied as supplementary videos.

To test whether CwhA activity leads to killing of *S. aureus*, bacterial growth was assessed on LB agar plates after treatment of bacteria with CwhA for different time periods (Figure 3C). 15 min after addition of CwhA, only 60% of initial CFU were recovered, while both PBS and lysozyme only led to a minor reduction (∼10 %). Treatment with CwhA led to further reduction of CFU over time, and after 120 min nearly all staphylococci were killed. In contrast, after the initial drop, CFU in the PBS control remained stable, and only 50% CFU reduction were achieved with lysozyme. To assess if bacterial killing was mediated by cell wall lysis, the effect of CwhA on *S. aureus* was analysed using an *S. aureus* strain expressing green fluorescent protein (GFP) in the cytoplasm. Lysis of bacterial cells should lead to leakage of cytoplasmic contents and thereby loss of fluorescence, which can be visualized by automated live cell imaging microscopy. While the fluorescence of *S. aureus* cells treated with PBS or BSA remained stable, the GFP signal rapidly decreased by 50 % within one hour in the CwhA-treated cells (Figure 3D; Supplementary Video S1). This indicates that CwhA indeed has a bacteriolytic effect on staphylococci, leading to the rupture of the bacteria and release of cytoplasmic content.

### CwhA produced by *A. fumigatus* lyses *S. aureus* in co-culture

To determine if the native CwhA protein produced by *A. fumigatus* shows the same lytic activity against *S. aureus* as demonstrated for the recombinant protein, we generated a mutant strain deficient in CwhA (Δ*cwhA*), as well as a mutant strain that overproduces the protein, using the constitutively active promoter of the *A. fumigatus gpdA* gene (P*gpdA*::*cwhA*). While no expression of *cwhA* was detectable in the deletion mutant, *A. fumigatus* P*gpdA*::*cwhA* showed highly increased *cwhA* gene expression compared to the wild type (Figure 4A). Western blot analyses using a monoclonal antibody directed against CwhA confirmed protein overexpression in *A. fumigatus* P*gpdA*::*cwhA* (Figure 4B), which was especially pronounced in cellular extracts. Secretion of the protein, however, was relatively weak during growth in *Aspergillus* minimal medium (AMM) (Figure 4B). We also confirmed our previous literature-based assumption that the protein is not produced at conventional laboratory *in vitro* culture conditions, as no protein band was detectable in protein extracts of the *A. fumigatus* wild type (Figure 4B). To determine the functionality of CwhA produced by *A. fumigatus*, the three strains were tested for their capacity to induce bacterial lysis in *S. aureus*. We observed a strong decrease in bacterial numbers after 24 hour co-incubation of bacteria with fungal mycelia of the P*gpdA*::*cwhA* strain (Figure 4C). In contrast, there were no differences in the bacterial counts after co-cultivation with fungal mycelia of the wild type and the Δ*cwhA* deletion strain. In conclusion, CwhA produced as native protein by *A. fumigatus* wild type shows antibacterial features similar to the recombinant protein, but the gene is only weakly expressed in axenic laboratory cultures.

**Figure 4.**
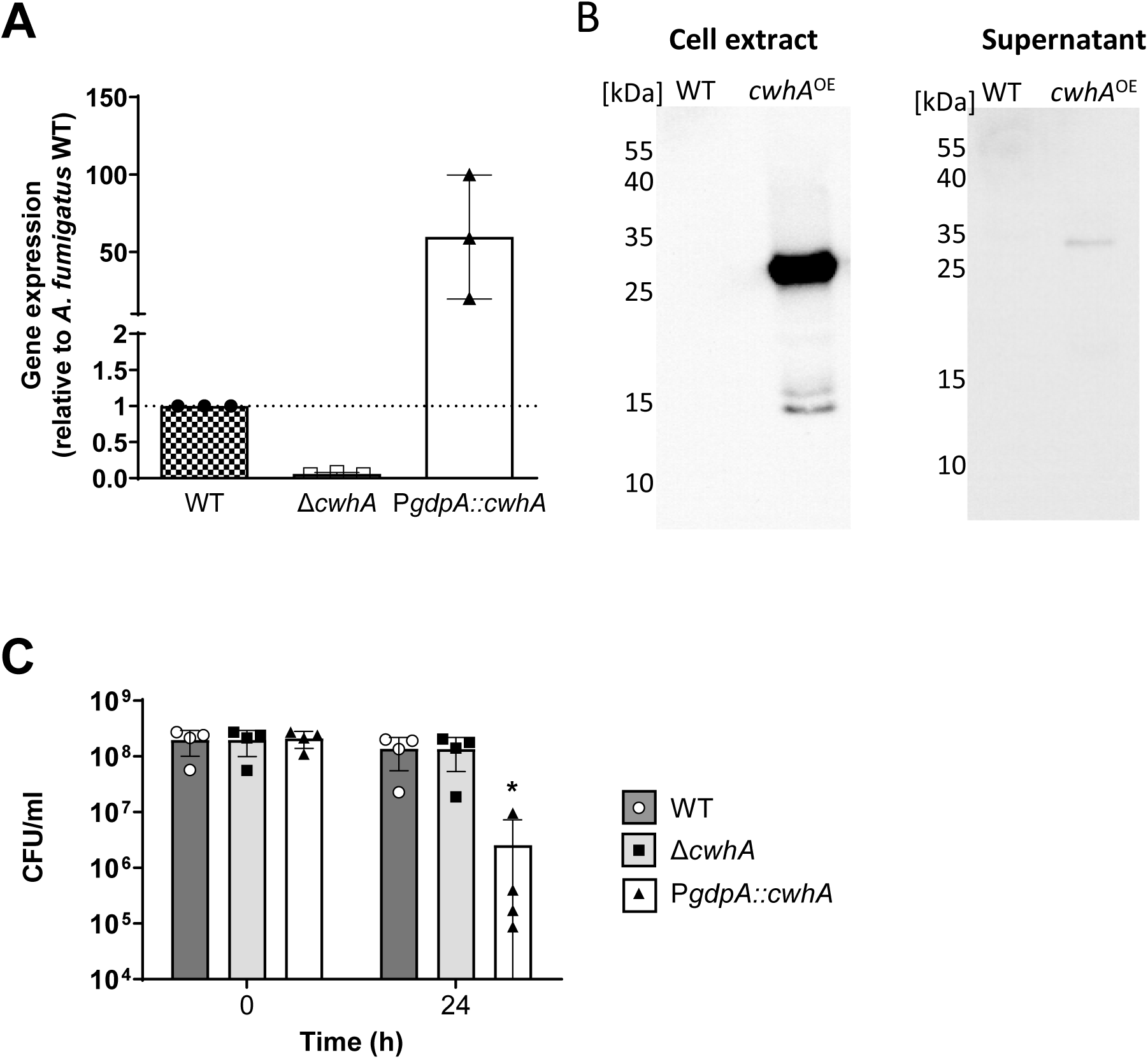
Over expression of *cwhA* in *A. fumigatus* increased bacterial lysis. (A) Expression of *cwhA* determined by quantitative RT-PCR of *Aspergillus fumigatus*. Relative expression of *cwhA* was determined by the 2-ΔΔCt method with *cox5* and *act1* as housekeeping genes; expression in *A. fumigatus* CEA17 Δ*akuB^KU80^*(WT) was set to 1. (B) CwhA protein expression was determined in whole cell extracts and culture supernatants of the *A. fumigatus* CEA17 Δ*akuB^KU80^*strain (WT) and its isogenic overexpressing mutant P*gpdA*::*cwhA* using a monoclonal antibody raised against recombinant CwhA protein. (C) Survival of *S. aureus* after 24 h co-incubation in Tris-Buffer with *A. fumigatus CEA17 ΔakuB^KU80^* as wild type and its isogenic *cwhA* mutants. Data from four independent experiments is shown as scatter plot with bars; bars represent mean and error bars SD. *Statistical analysis by one-way ANOVA and Tukey’s multiple comparisons test showed a significant difference in CFU after 24 h coincubation with *A. fumigatus* P*gpdA::cwhA* compared to either the WT (p=0.046) or the deletion strain Δ*cwhA* (p=0.048).

### CwhA gene expression in *A. fumigatus* is induced by the presence of some Gram-positive bacteria

CwhA was identified as one of the most abundant fungal proteins in BAL samples in mice with invasive aspergillosis (Machata *et al*., 2020). Furthermore, a 5-fold upregulation of the expression of Afu8g00360, the gene encoding CwhA in *A. fumigatus* Af293, was shown in a previous study by *in vivo* transcriptome analysis of BAL samples from infected mice (McDonagh *et al*., 2008). To identify possible factors that lead to the induction of CwhA *in vivo,* we analysed its gene expression under two conditions that mimic environmental factors in the lung during IPA. First, we tested if oxygen availability has an impact on *cwhA* expression, as hypoxic conditions occur in infected lungs (Grahl *et al*, 2011; Hillmann *et al*, 2015b). Our qRT-PCR data revealed no significant difference in the transcript levels of CwhA between cultures exposed to normoxia (21 % O_2_) and hypoxia (0.2 % O_2_; expression 1.067 ± 0.48 compared to normoxia). Secondly, expression in the presence of host cells was analysed. Infection of the human alveolar epithelial cell line A549 for 6 h did not lead to an increase of *cwhA* expression (Figure 5B). However, a 5-fold increase in *cwhA* expression was observed when the infection period was extended to 24 h, at which host cells were partially disrupted by invading fungal mycelia. These data show that reduced oxygen availability has no impact on the *cwhA* gene induction, and that the sole presence of host cells is insufficient to promote gene expression. Increased expression of *cwhA* could only be observed at a late stage of infection, which may be attributable to a response to factors released from damaged host cells, or be merely the result of nutritional starvation.

**Figure 5.**
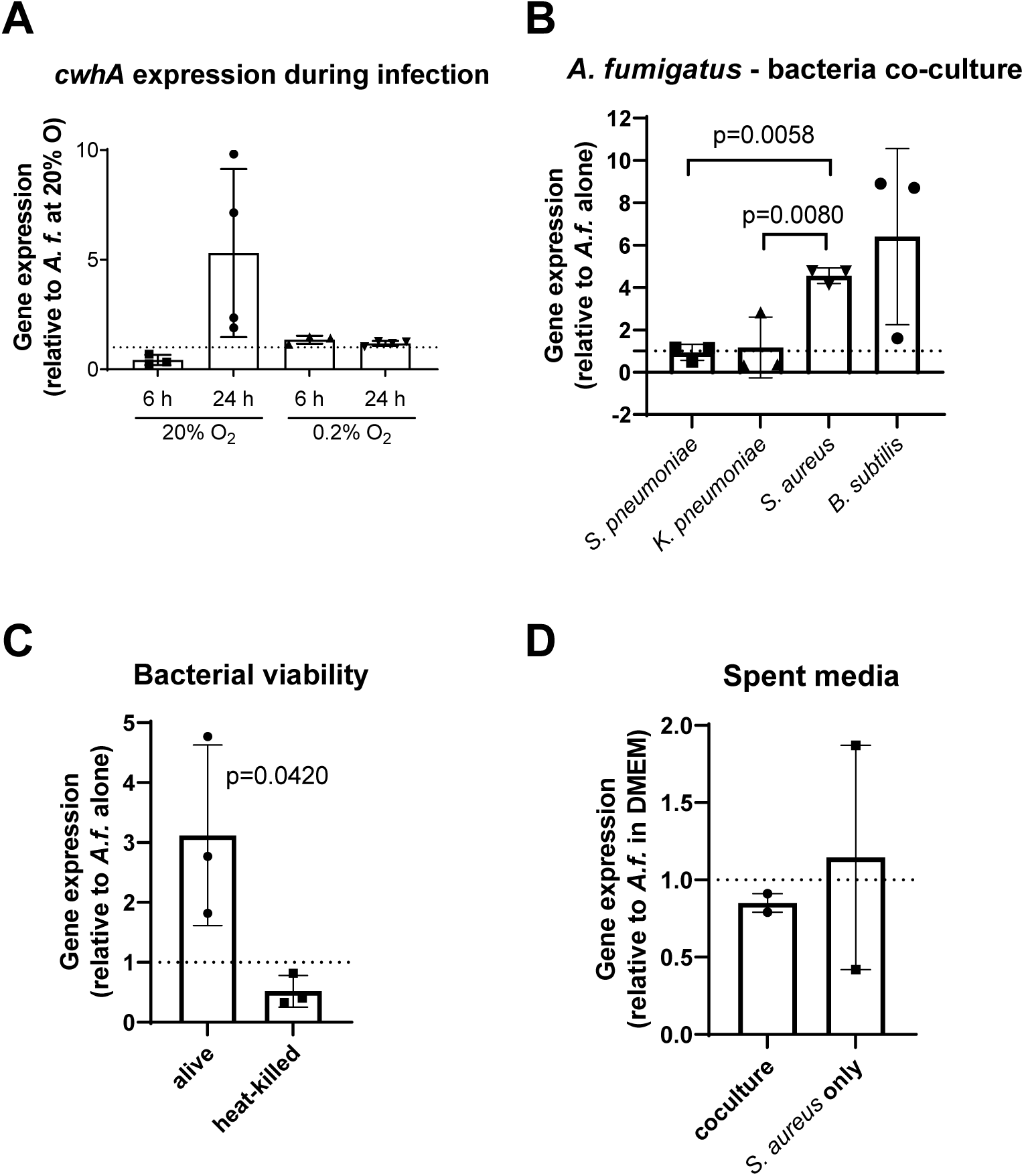
Impact of oxygen, host cells, and bacteria on *cwhA* expression. *cwhA* expression was determined by qRT-PCR using RNA isolated from *A. fumigatus* mycelia (A) during *in vitro* infection of A549 cells at 20 % and 0.2 % oxygen, (B) from co-cultures in DMEM with *B. subtilis*, *S. pneumoniae*, *K. pneumoniae* and *S. aureus*, respectively, (C) in the presence of living or heat-killed *S. aureus* (Newman), and (E) grown in spent media (sterile-filtered culture supernatant of *S. aureus* Newman) alone or in co-culture with *A. fumigatus* for 4 h in DMEM. Data is represented as scatter plot with bar (A-C: mean ±SDs, three independent experiments; D: mean and range, two independent experiments). Data was analysed by ordinary one-way ANOVA and Tukey‘s multiple comparison test (A, B), or two-tailed unpaired t-test (C), p values < 0.05 are indicated in the graphs.

As we found CwhA to be effective against some Gram-positive bacteria, we next assessed whether fungal proximity to bacteria promotes expression of *cwhA* (AFUB_086210). Co-incubation of *A. fumigatus* with either *S. aureus* and *B. subtilis* led to a clear increase in *cwhA* expression (4-and 6-fold, respectively), while the expression remained unaffected by co-incubation with *S. pneumoniae* or *K. pneumoniae* (Figure 5C). The induction was dependent on bacterial viability as determined with heat-inactivated *S. aureus* (Figure 5D). Thus, the mere presence of bacterial surface structures does not seem to be sufficient for inducing *cwhA* expression. We also tested if soluble compounds released by bacteria, or changes in the culture environment due to bacterial metabolism, stimulate the expression of *cwhA*. Therefore, fungal mycelia were incubated in sterile-filtered supernatants of bacterial fungal-bacterial co-cultures (Figure 5E). Neither type of supernatant led to increased *cwhA* expression, indicating that soluble factors and nutritional starvation alone are not the sole inducing factors, and that a combination of bacterial activity and physical interaction might be required.

### Recombinant CwhA causes no damage to amoeba, host cells, and fungal cells

We next assessed the role of CwhA in interaction with organisms other than bacteria that can occur in close proximity to *A. fumigatus* in nature and might be affected by the secreted protein. Amoebae are commonly found in damp soil where they co-localize with *A. fumigatus* (Hillmann *et al*., 2015a). We therefore tested the susceptibility of the amoeba *Dictyostelium discoideum* to CwhA. At early time points, the recombinant protein had a minor growth-promoting effect, whereas after 26 h a slight reduction in numbers of amoebae was observed (Figure 6A). Since *A. fumigatus* CwhA was produced in detectable amounts in lung tissue during murine IPA, we next determined if CwhA contributes to host cell damage. CwhA concentrations ranging from 0.4 to 25 µg/ml did not cause detectable necrotic cell damage of murine alveolar macrophages (MH-S) and human pulmonary epithelial cells (A549) within 12 h of exposure (Figure 6B). Last, we also analysed if CwhA has an inhibitory effect on *A. fumigatus* grown as planktonic culture or as biofilm. Addition of recombinant CwhA to *A. fumigatus* in AMM had a stimulatory impact, rather than an inhibitory one, with its effects on fungal growth comparable to or slightly more pronounced than the control protein bovine serum albumin (BSA) (Figure 6C, D). This likely indicates that CwhA is utilized as an additional nutrient source in these settings. In summary, recombinant CwhA exhibits lytic activity against some Gram-positive bacteria, but not against the Gram-negative bacteria tested, amoebae, host cells, and *A. fumigatus* itself.

**Figure 6.**
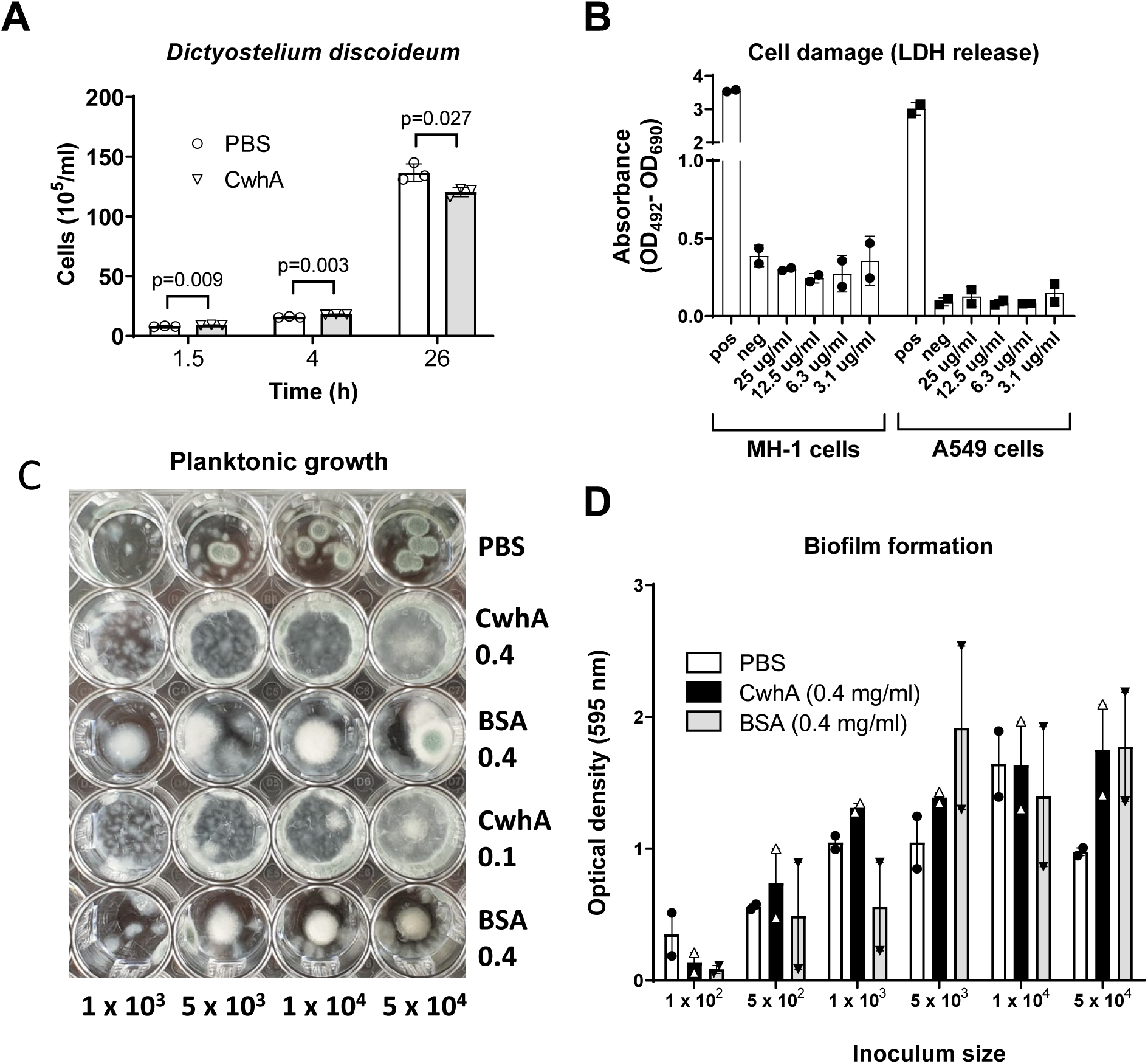
Effect of CwhA on amoeba, host cells and fungi. (A) Cell numbers of *Dictyostelium discoideum* in HL5 media following different incubation times with 130 µg/ml CwhA or PBS. Data from three individual experiments is shown as column bars with mean ±SD, symbols represent the individual data points. Data was analyzed by 2way ANOVA (no significant impact of treatment) and per time point by two-tailed unpaired t-test (p values indicated in the graph). (B) Cell damage determined by lactate dehydrogenase (LDH) release of murine alveolar macrophages (MH-S) and human pulmonary epithelial cells (A549) after treatment with CwhA (3.1 - 25μg/ml). (C) Effect of CwhA on *A. fumigatus* CEA10 during planktonic growth or (D) on biofilm formation. *A. fumigatus* spores (1 x 10^2^/ml to 5 x 10^4^/ml) were inoculated in AMM supplemented with PBS, CwhA or Bovine serum albumine (BSA) at 0.4 and 0.1 mg/ml and incubated at 37°C for 48h either shaking (300 rpm)(C) or non-shaking (D) for adherent growth.

### CwhA acts as an endopeptidase on peptidoglycan of Gram-positive bacteria containing L-lysine in the peptidoglycan stem peptide

The observation that recombinant CwhA was only active against certain bacteria suggested substrate specificity. Although both *S. aureus* and *B. subtilis* are Gram-positive, they differ in their PG composition (a detailed description is provided with the supplementary material). The most important difference is on position three of the stem peptide, where *S. aureus* harbours L-lysine, which is typical for Gram-positives. *B. subtilis* contains meso-diaminopimelic acid (mDpm) instead, normally found in the PG of Gram-negatives (Atrih *et al*, 1999; Do *et al*, 2020; Kim *et al*, 2015; Vollmer *et al*, 2008). To determine the exact hydrolytic specificity of CwhA, PG isolated from *S. aureus* and *B. subtilis* was treated with the muraminidase mutanolysin alone or with a combination of mutanolysin and CwhA. The cleavage products were separated by UHPLC and analysed by MS. When the digestion was performed with mutanolysin alone, both strains showed the expected muropeptides (Figure 7A-I, B-I) (Atrih *et al*., 1999; Do *et al*., 2020). For *S. aureus,* these were mostly disaccharides (DS) with a stem peptide of five amino acids (pentapeptide), often harbouring a pentaglycine bridge. Several stem peptides can be cross-linked indirectly *via* these pentaglycine bridges, resulting in multimeric muropeptides (Figure 7A-I and supplementary Figure 3). When *S. aureus* PG was treated with CwhA alone, the enzyme hydrolysed a bond within the stem peptide, thereby segregating most of the peptide part of PG from the glycan strands (Figure 7A-II). The resulting masses matched with peptide residues of four to seven subunits cross-linked *via* the pentaglycine bridge. The *m/z* values observed in the other peaks belonged to glycan strands of one to nine DS-units that still contained one dipeptide (L-alanine -amidated D-glutamate) per N-acetylmuramic acid (NAM) residue. Therefore, the cleavage site of CwhA must be located between the second and the third amino acid of the stem peptide (in *S. aureus* this is amidated D-glutamate and L-lysine, respectively). When digested with both enzymes, mutanolysin and CwhA, the entire macromolecule was degraded into DS-dipeptides, the smallest PG cleavage product (Figure 7A-III and A-IV). Three different variants of DS-dipeptides were observed: i) unmodified DS-dipeptide is the most abundant form, ii) DS-dipeptide without N-acetyl glucosamine (NAG), and iii) DS-dipeptide with an additional O-acetylation known to occur at position six of NAM ^39,40^ (Figure 7A-III, a-c). The unmodified DS-dipeptide was also found in PG treated with mutanolysin alone (Figure 7 A-I), but its amount increased by a factor of 165 when CwhA was added (Figure 7A-III, A-IV).

**Figure 7.**
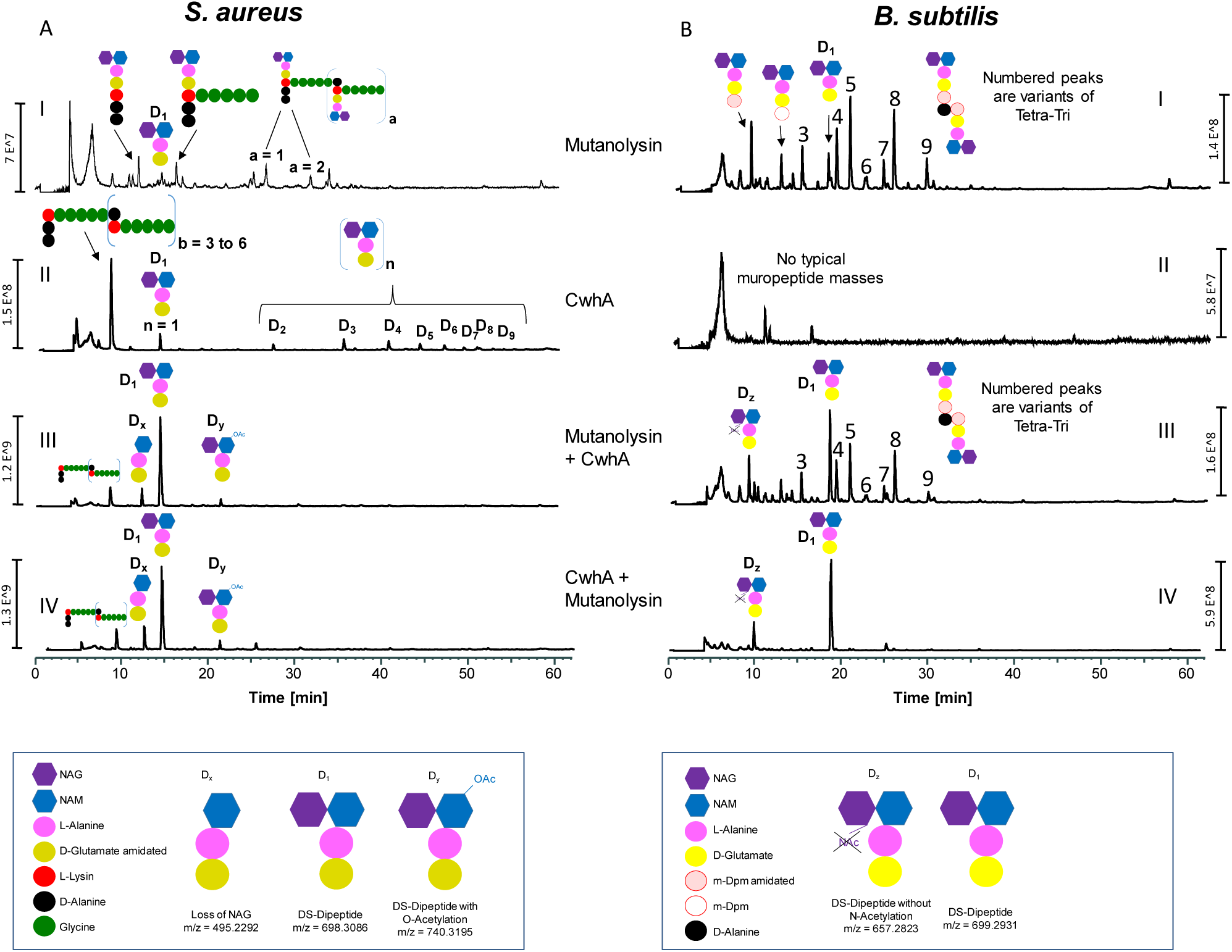
Muropeptide profile of *S. aureus* SA113 and *B. subtilis* (Ehrenberg 1835) obtained by UPLC/MS after treatment with mutanolysin or/and CwhA. Masses of peaks are shown in Supplementary Table 3. Numbered peaks are variants of Tetra-Tri.

In contrast to *S. aureus*, *B. subtilis* contains mDpm at position three of the stem peptide. Within mature PG, the stem peptides are shortened to three or four amino acids by native enzymes, resulting in the formation of tetra-and tripeptides. Most muropeptides released by mutanolysin were variants of tetra-tri-dimers with various chemical modifications, such as amidation of mDAP and N-deacetylation of NAM (Figure 7B-I) (Atrih *et al*., 1999). Treatment of *B. subtilis* PG with CwhA alone did not release any detectable muropeptides (Figure 7 B-II). However, when CwhA was added to PG that had been pre-treated with mutanolysin, the amount of DS-dipeptide increased while, concomitantly, the other muropeptides decreased (Fig. 6, compare panel B-III with panel B-I). When PG was initially treated with CwhA and subsequently with mutanolysin (resulting *de facto* in a prolonged CwhA treatment), even *B. subtilis* PG was completely digested into DS-dipeptides, as shown in Fig. 7, panel B-IV. Therefore, CwhA also accepts amidated mDpm-containing PG as substrate, not only the one with L-lysine. Unlike for *S. aureus*, the released peptide part was not detectable for *B. subtilis*. For the most part, in *B. subtilis* PG only two adjacent stem peptides are cross-linked without an interpeptide bridge in contrast to three to seven stem peptides in *S. aureus*. This results in molecules of a very small size that does not facilitate retention on the column used.

Taken together our data indicates that the enzyme CwhA is a fungal endopeptidase hydrolysing the peptide bond between amino acids two and three of the PG stem peptides. It thereby seems to prefer L-lysine over mDpm on position three. However, with prolonged treatment, even PG containing amidated mDpm was degraded into DS-dipeptides, potentially explaining the lytic band detected in the zymogram gels (Figure 2B). Further cleavage experiments demonstrated that CwhA in combination with mutanolysin efficiently degraded PG of two other Gram-positive bacterial strains containing L-lysine instead of mDpm, giving rise to a major peak for the DS-dipeptide (Supplementary Figure 4). One of the tested bacteria was an MRSA strain with the same PG structure as *S. aureus* Newman, the other one was *S. pneumoniae* (Klein 1884) that contains alanyl-serine cross-bridges or is directly cross-linked (Garcia-Bustos *et al*, 1987; Schleifer & Kandler, 1972). *B. subtilis* was re-tested and showed a similar muropeptide profile as PG treated with mutanolysin alone, and therefore no activity of CwhA could be detected, most likely because the incubation time was shorter than when mutanolysin was added after treatment with CwhA. These data suggest that CwhA is a broad-spectrum enzyme that efficiently degrades the PG of various Gram-positive bacteria containing L-lysine at position 3 of the stem peptide, leading to high accumulation of the smallest PG cleavage product, DS-dipeptide. Thus, variations in the stem peptides explain the observed differences in CwhA efficacy against different bacterial species.

### CwhA-mediated PG cleavage induces cytokine responses

While CwhA itself did not damage host cells (Figure 6B) and deletion of *cwhA* did not affect fungal survival if challenged with human macrophages (Supplementary Figure 5), we hypothesized that CwhA activity might indirectly impact immune cells by processing of bacterial cell walls leading to release of ligands for pattern recognition receptors. Gresnigt et al. showed that NOD2 receptor deficiency renders mice more resistant to IPA, and that risk patients with NOD2 polymorphisms are less predisposed to IP (Gresnigt *et al*, 2018). Bacterial Muramyl-dipeptide (MDP) is a known NOD2 agonist and can synergistically increase *Aspergillus-*induced cytokine levels (Li *et al*, 2012; Zhang *et al*, 2008). Additionally, NOD2 stimulation with its MDP in the presence of *A. fumigatus* was shown to positively regulate cytokine production and to decrease phagocytosis and fungal killing (Gresnigt *et al*., 2018). NOD2 has also been shown to be involved in chitin-induced anti-inflammatory signalling (Wagener *et al*, 2014); however, it still remains unclear how NOD2 is activated by *A. fumigatus*. We hypothesized that CwhA-mediated cleavage of PG into muropeptides that serve as NOD2 ligands could contribute to the effects of IPA in an *in vivo* setting where Gram-positive bacteria are present. We therefore measured IL-1α and TNF-α in supernatants of human PBMCs after stimulation. We confirmed cytokine induction by a combination of MDP and heat-killed conidia, and furthermore observed a significant increase of both cytokines after stimulation with CwhA-degraded PG (Figure 8). Thus, *in vivo* bacterial cleavage by CwhA might affect immune responses and IPA severity.

**Figure 8:**
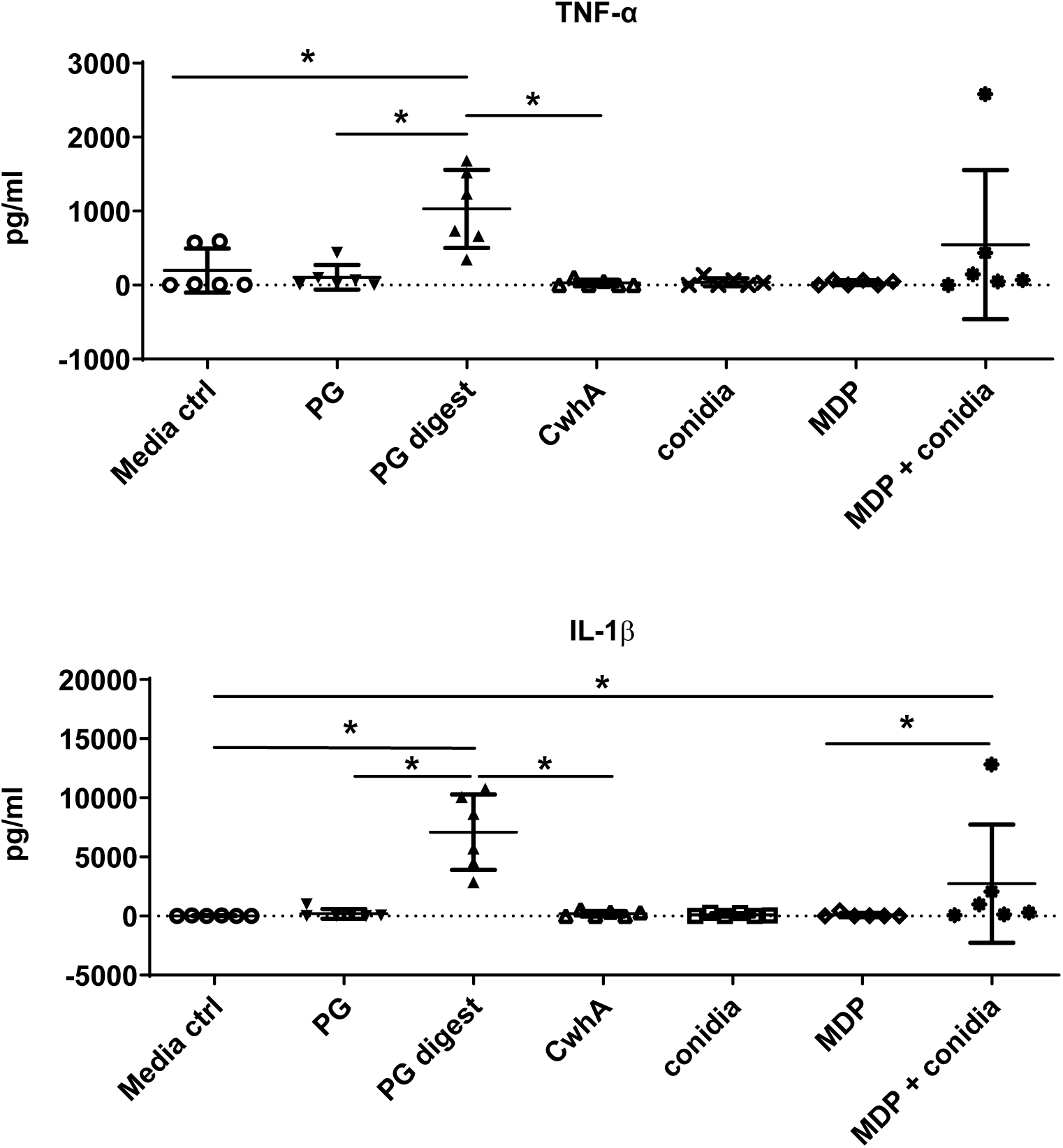
PG cleaved by CwhA leads to cytokine induction of human PBMCs. *Aspergillus*-MDP-induced TNF-α and IL-1β levels in culture supernatants of PMBCs (5x 10^5^) of healthy volunteers (n=6) presented as scatter dot plot with mean ± SD. The data was analyzed by pairwise comparison using the Wilcoxon matched-pairs signed rank test. Asterisks indicate significant differences (exact two-tailed p value of 0.0313). Media ctrl: Medium (RPMI 1640^+^ with 10% human serum) only; PG: *S. aureus* PG only; PG digest: *S. aureus* PG + 20 µg CwhA; CwhA: 20 µg CwhA; conidia: 1 x 10^7^/ml heat-killed conidia of *A. fumigatus* CEA10; MDP: 10 µg/ml MDP; MDP + conidia: 10 µg/ml MDP and 1 x 10^7^/ml heat-killed conidia of *A. fumigatus* CEA10.

## Discussion

As an environmental, saprophytic fungus, *A. fumigatus* inhabits ecological niches with high microbial diversity and cell density leading to a strong interspecies communication. It is thus not surprising that in addition to metabolic versatility antimicrobial mechanisms evolved in fungi to promote survival in highly competitive environments (Macheleidt *et al*, 2016; Scharf *et al*., 2016). For example, filamentous fungi produce an enormous structural diversity of secondary metabolites with a broad spectrum of activities and functions, *e.g*., for communication with other microorganisms or combating bacterial competitors (Macheleidt *et al*., 2016).

Here, we found a special protein that has similar functions. It is the previously uncharacterized fungal protein B0YAY0, a secreted endopeptidase that specifically cleaves distinct bonds in the stem peptide of bacterial PG. Due to its activity, and similarity of the functional domains with bacterial cell wall hydrolases, we renamed the protein cell wall hydrolase A (CwhA). Bacterial cell wall hydrolases are essential for cell division, cell wall recycling, and differentiation. Therefore, bacteria produce a wide range of these enzymes that differ in PG-binding and cleavage sites and their substrate specificity (Do *et al*., 2020). Some murein hydrolases, so-called bacteriolysins, are also used by bacteria to eliminate other bacterial strains that share the same ecological niche and compete for nutrients and other limiting factors (Wittekind & Schuch, 2016). A well-characterized example is lysostaphin, which is produced by *Staphylococcus simulans* biovar *staphylolyticus.* This metalloenzyme cleaves pentaglycine cross-bridges in PG of other *Staphylococcus* strains to eliminate competitors (Bastos *et al*, 2010). Cell wall hydrolases are also produced by animals, including humans: Lysozyme, which we used in our study as a positive control, is highly abundant in various body fluids and involved in host defence against bacteria by cleavage of peptidoglycan and subsequent bacterial lysis (Ragland & Criss, 2017). The conservation of functional lysozymes throughout the animal kingdom underscores their value in combating bacteria (Callewaert & Michiels, 2010); it is thus not surprising that fungi like *A. fumigatus* produce proteins with similar functions to defend the habitat against bacteria.

CwhA is not produced under standard laboratory culture conditions, which explains why its function was not discovered earlier. The transcription of *cwhA* was, however, upregulated during co-culture with some bacteria. Physical contact with bacteria also induces production of secondary metabolites with antibacterial potential (Fischer *et al*., 2018; Netzker *et al*., 2015; Schroeckh *et al*., 2009). This suggests a general fungal strategy to produce antimicrobial metabolites and proteins only in the presence of other microorganisms when these compounds are beneficial. Of note, although increased *cwhA* transcription during co-culture with *S. aureus* was observed, the level of induction was only moderate. Likely, the transcriptional upregulation observed does not lead to sufficient production of secreted CwhA to kill a substantial number of bacteria in the assays employed here, thereby explaining why only the overexpression strain but not *A. fumigatus* WT showed substantial bacterial killing in the co-cultivation experiments. This is in contrast to the local induction of the secreted protein *in vivo* during IPA in mice. Although we did not observe an upregulation of *cwhA* under hypoxia or in contact with lung epithelial cells, it is possible that a combination of stimuli drives expression *in vivo*. In *Aspergillus fumigatus* Af293 *cwhA* (Afu800360) is located in a “supercluster” (Afu8g00100 – Afu8g00720) on chromosome 8. This supercluster region includes gene clusters responsible for the biosynthesis of fumitremorgin, fumagillin and pseurotin A (Wiemann *et al*, 2013). The regulation of genes in this chromosomal region is rather complex and includes a number of different transcription factors (Dhingra *et al*, 2013; Wiemann *et al*., 2013; Yu *et al*, 2018). For CwhA such a regulator has not been identified yet that explains its apparently higher expression *in vivo* (mouse lungs) than *in vitro* in mono culture of *A. fumigatus*. Furthermore, we also identified two other uncharacterized fungal proteins, B0Y269 and B0Y9E0, which were highly abundant in BAL samples of mice with invasive aspergillosis, and shared common motifs with CwhA (Machata *et al*., 2020). Like CwhA, these proteins appear not to be produced *in vitro* but the gene encoding B0Y269 was highly upregulated *in vivo* (McDonagh *et al*., 2008). These common features with CwhA are a hint that the proteins might have similar functional roles and could work in concerted action with CwhA, possibly showing different substrate specificities, with preference towards other bacteria. Further work needs to be done to investigate these proteins in more detail and to elucidate theirs functions for fungal physiology

Since we found no indication that CwhA is cytotoxic or that *cwhA* is important for the interaction with macrophages, it seems unlikely that CwhA directly affects virulence. CwhA might, however, influence interactions between *A. fumigatus* and bacteria during polymicrobial pulmonary colonisation/infection, for example in CF patients. The role of allergic bronchopulmonary aspergillosis in CF patients has been firmly established (Carsin *et al*, 2017), but the effects of colonisation without infection on CF remain unclear. By reducing *S. aureus* colonisation, *A. fumigatus* might promote growth of Gram-negative bacteria such as *P. aeruginosa* that are not impacted by CwhA, and thereby contribute to disease development.

Another effect that accompanies CwhA-mediated bacteriolysis is the accumulation of increased levels of muramyl dipeptides (MDP) due to the cleavage of bacterial peptidoglycan. Since NOD2 is the sensing protein for MDP in host cells (Girardin *et al*, 2003), we tested the CwhA cleavage products for immune stimulation of human PBMCs. The increased cytokine levels upon treatment with PG pre-digested beforehand with CwhA confirmed our hypothesis that the release of CwhA during infection could affect the immune stimulation in the host. This finding is particularly intriguing from the immunological point of view, as Gresnigt et al. showed that NOD2 stimulation by a combination of *A. fumigatus* conidia and MDP suppresses fungal killing in human monocyte-derived macrophages (MDMs) or monocytes and leads to increased cytokine production (Gresnigt *et al*., 2018). The production and activity of CwhA by *A. fumigatus* in the lung of a host who harbors microbiota and is constantly exposed to external bacteria could explain the MDP-mediated stimulation and thus the relevance and importance of the NOD2 receptor for *Aspergillus* infection. Interestingly, the positive impact of cell wall hydrolases on the immune tolerance toward pathogenic bacteria was already previously described (Rangan *et al*, 2016). The tolerance to enteric bacteria was reported to be increased upon release of an NlpC/p60 PG hydrolase by *E. faecium* in *C. elegans* through *tol-1* signaling (Rangan *et al*., 2016). Further studies are necessary to get a better insight into the validity and mechanism of the CwhA contribution to host immune responses and fungal survival in the host.

In summary, CwhA is the first secreted antibacterial protein identified in *A. fumigatus*. As an endopeptidase it cleaves PG of Gram-positive bacteria, leading to bacterial lysis and the release of PG cleavage products that stimulate cytokine production of human immune cells. Its expression in lungs of mice infected with *A. fumigatus* can be partially explained by the presence of bacteria, but is likely triggered by complex stimuli. Together with other fungal factors that influence bacterial growth, it might impact the microbiome in both environmental and clinical niches inhabited by *A. fumigatus*.

## Methods

### Lung biopsy and MALDI-IMS

Murine lungs were obtained from immunocompromised animals infected intranasally with *A. fumigatus* as part of a previous study (Machata *et al*., 2020). Lungs were stored in 4 % formalin (Histofix), samples were embedded in paraffin, cut into 5 µm sections, and transferred onto conductive ITO slides (Bruker Daltonics, Bremen, Germany). For correlation of the peptide mass, 1 µl of 200 ng/ml recombinant CwhA was spotted next to the tissue sections. The sample preparation was performed as described previously (Hoffmann *et al*, 2019). Shortly, the sections underwent deparaffinization, rehydration, pH-conditioning, and antigen retrieval. For trypsin deposition, the SunCollect (SunChrom GmbH, Frankfurt, Germany) was used. The tryptic digestion was performed using the digestion chamber SunDigest (Sunchrom GmbH) in “smart mode”, using the following protocol: number of steps: 2; step 1: 900 s, 0 % fan speed, base temperature: 50 °C, cover temperature: 45 °C; step 2: 6300 s, 2 % fan speed, base temperature: 50 °C, cover temperature: 45 °C. The matrix used for all sections consisted of 30 mg/ml 2,5-dihydroxybenzoic acid (DHB) in 50 % acetonitrile and 0.2 % trifluoroacetic acid. Matrix application was performed with the SunCollect with the following parameters: five layers; flow rate for layer 1: 10 µl/min; flow rate for layers 2 to 5: 35 µl/min; speed x: low 3, speed y: medium 1, vial distance: 2 mm, Z position: 29 mm. MALDI-TOF – IMS data acquisition was performed using the UltrafleXtreme mass spectrometer (Bruker Daltonics). The measurements were carried out in positive ion reflective mode, with 50 µm pixel size, medium laser beam size, 200 shots per position, in a mass range of *m/z* 500 to 4000. The Bruker peptide calibration standard was used for external mass spectrometer calibration. After MALDI-IMS measurements, the sections were washed twice with 80 % ethanol and PAS (periodic acid-Schiff) -stained for histopathological annotation. The slides were scanned using the Panoramic DESK (Sysmex, Germany). For data and imaging analysis, the SCiLS Lab software, version 2020a Premium 3D (Bruker Daltonics) was used. All data were TIC-normalized. For *in silico* digestion the ProteinProspector MS-Digest tool was used (http://prospector.ucsf.edu).

### Microorganisms, media and cultivation

*A. fumigatus* wild-type strains CEA10 (FGSC A1163; wild type used for most experiments unless otherwise stated) and its derivatives CEA17 Δ*akuB^KU80^* (da Silva Ferreira et al, 2006), CEA17 Δ*akuB^KU80^*Δ*cwhA* (this study), and CEA17 Δ*akuB^KU80^*::P*gpdA*::*cwhA* (this study) were used for *in vitro* experiments. The bioluminescent *A. fumigatus* strain C3 (based on CGS144.89, the wild type origin of CEA10) was kindly provided by Matthias Brock (University of Nottingham, UK) and used for murine infection experiments in a previous study (Brock *et al*, 2008) which provided the lung tissue used here. Fungal spores were grown at 37°C on *Aspergillus* minimal medium (AMM) agar for three days (Brakhage & Van den Brulle, 1995). When required, the medium was supplemented with 1 µg/ml pyrithiamine hydrobromide (Sigma Aldrich). Conidia were harvested from agar plates with water and counted using the CASY Cell Counter (Roche Innovatis). Fungal mycelium was harvested with Miracloth (Merck Millipore) from liquid cultures of *A. fumigatus* strains grown overnight in AMM medium at 37°C, after washing with water or with sterile PBS when required. *Staphylococcus aureus* (Newman and SA113), *Bacillus subtilis* (DSM3256), *Streptococcus pneumoniae* (ATCC 6303), *S. epidermidis* (IMET10845)*, Klebsiella pneumoniae* (ATCC 700603)*, Pseudomonas aeruginosa* (PAO1), and *Enterococcus faecalis* (clinical blood isolate BK014967, kindly provided by Steffen Höring, University Hospital Jena) wild type strains were used for peptidoglycan isolation, lysis assays and/or co-cultivation experiments, culture details are provided below. Additional *S. aureus* strains (multiresistant, MRSA) used to test the effect of CwhA are listed in Supplementary Table 4. The GFP-expressing *Staphylococcus aureus* strain (6850/pALC1743) (Balwit *et al*, 1994; Kahl *et al*, 2000) was used to determine lytic activity of recombinant CwhA. Bacterial cultures were grown in LB medium or on LB agar plates. In survival assays after co-cultivation with *A. fumigatus,* 0.1 mg/ml nystatin (Merck) was added to LB agar plates to prevent fungal growth.

### Genetic manipulations and purification of recombinant protein

A list of oligonucleotides used in this study is provided with the supplementary material (Supplementary Table 5). The generation of PCR fragments for all genetic manipulations was carried out using the Phusion High Fidelity DNA Polymerase (NEB). *A. fumigatus* deletion and overexpression strains were generated using CEA17 Δ*akuB^KU80^*as the parental strain. The AFUB_086210 (*cwhA*) deletion (resulting in strain CEA17 Δ*akuB^KU80^*Δ*cwhA*) was performed by replacing the gene with a pyrithiamine resistance cassette generated with ptrA_for_II and ptrA_rev_II primers from plasmid pSK275 (Szewczyk *et al*, 2006) using 5’ and 3’ flanking regions of AFUB_086210, generated with primers Del_p60A_F1 and Del_p60A_R2, and Del_p60A_F3 and Del_p60A_R4, respectively. The deletion was achieved by homologous recombination following transformation of protoplasts as previously described (Weidner *et al*, 1998). Correct deletion was confirmed by PCR and by Southern Blot: The gene-specific DNA probe was generated by PCR using the primers Del_p60A_F1 and Del_p60A_R4. Genomic DNA isolated from the wild type and the mutant strains, was digested with the restriction enzymes XhoI and SacI (NEB). The overexpressing strain CEA17 Δ*akuB^KU80^*::P*gpdA*::*cwhA* was constructed using the vector pSK379 (Szewczyk & Krappmann, 2010) (kindly provided by Sven Krappmann, University of Erlangen) containing a pyrithiamine cassette and the P*gpdA* promoter. The *cwhA* gene was amplified with primers Expr_p60A_for and Expr_p60A_rev, using genomic DNA derived from *A. fumigatus* CEA10 as template, and integrated into the plasmid linearized with *Pme*I (NEB) restriction enzyme. The CEA17 Δ*akuB^KU80^*::P*gpdA*::*cwhA* was obtained by homologous recombination following the transformation of protoplasts as previously (Weidner *et al*., 1998). For generation of the recombinant CwhA protein, the AFUB_086210 gene was amplified from genomic DNA of *A. fumigatus* CEA10 using o_p60AFor and o_p60ARev primers. The amplified DNA was cloned into the plasmid pRSETb (Addgene) at its multiple cloning site C-terminal of a His-tag sequence and transformed into *E. coli* DH5α (Invitrogen). Isolated plasmid from positive transformants was verified by sequencing and transformed into the *E. coli* KRX strain (Promega) for protein expression. Expression, induced by 0.1 % rhamnose, was continued at 30°C for 6 h. Cells were collected by centrifugation at 4°C, 10,000 x g for 45 min and lysed by French press. Protein purification was performed using immobilized metal chromatography (His-Trap, GE) and the Resource S ion exchange chromatography (Resource S, GE) on the ÄKTA (GE Health Life Sciences) purification system. The protein was concentrated using Amicon centrifugal units with 10kDa pore size (Millipore), the final protein concentration was evaluated by Bradford assay and protein purity was assessed by SDS-PAGE. Protein with >90 % purity was kept at 4°C for short term storage or supplemented with 10 % glycerol at -20°C for long term storage.

### Peptidoglycan isolation for LC-MS analysis

Peptidoglycan was isolated from *Staphylococcus aureus*, MRSA 181, *Bacillus subtilis* and *Streptococcus pneumoniae* overnight cultures as previously described in detail (Kühner *et al*, 2014). Briefly, cells from 3 ml overnight cultures grown in LB medium were harvested by centrifugation at 4°C and 10,000 x g for 10 min. Cells were resuspended in 1ml 1 M sodium chloride and boiled at 100 °C for 20 min in a heating block (ThermoStat plus, Eppendorf). The suspension was centrifuged at 10,000 x g for 5 min, cells were washed 4 times in ddH_2_0 and pellet was resuspended in 1 ml ddH_2_0. The sample was transferred to a water bath sonicator (Sonorex, Bandelin) for 30 minutes, after which 500 µl of 0.1 M Tris/HCl, pH 6.8 containing 15 µg/ml DNase and 60 µg/ml RNase were added. The suspension was incubated for 60 minutes at 37 °C with agitation at 180 rpm, 500 µl of a 50 g/ml trypsin solution was added and the suspension was incubated for another 60 minutes under the same conditions. Enzymes were inactivated by boiling the suspension for 3 minutes at 100 °C, cells were spun down at 10000 x g for 5 min and washed once in 1 ml ddH_2_0. The pellet was resuspended in 500 µl 1 M HCl and incubated for 4 h at 37 °C at 180 rpm. The suspension was centrifuged and washed with ddH_2_0 until the pH increased to 5-6. For digestion, the isolated PG was centrifuged, resuspended in 12.5 mM sodium dihydrogen phosphate (pH 5.5) and the optical density at 600 nm (OD_600_) was determined. Cell wall material was diluted to OD_600_ of 3.0 in the same buffer to a final volume of 200 µl, and 20 µl mutanolysin (5000 U/ml; Sigma-Aldrich) and/or 10 µl CwhA (4 mg/ml) or PBS were added. The digest was incubated at 37 °C, agitated at 150 rpm for 16 h, and the reaction was inactivated by boiling for 3 min at 100 °C. Samples were cooled on ice until the cleavage products were analysed by LC-MS.

### LC-MS analysis of cleavage products

Immediately prior to the analysis, muropeptides were reduced using sodium borohydrate (Sigma-Aldrich) and pH was adjusted to 3 by formic acid. Subsequent UHPLC analysis was performed as previously described ^68^, with instrument modifications. Muropeptide separation was perfomed with a WatersACQUITY UHPLC column (CSH C18 1.7 µm, 2.1x150 mm), from Waters with a guard column on an Agilent 1290 and a G42220 Bin Pump,baryp running in UHPLC mode with a maximum pressure limit of 1200 bar. Injection volume was 15 µl. The mobile phase consisted of LC-MS grade water (VWR) with 0.2 % formic acid (Sigma-Aldrich) A) and LC-MS grade methanol (VWR) with 0.2 % formic acid (B). The flow rate was 0.265 ml/min. For the initial 4 min, the mobile phase was held at 100 % A and the flow was discarded, in order to eliminate salt contaminants. From 4.10 min the gradient started with 3 % B and continued to 30 % B at 57 min. The column was washed for 5 min with 30 % B, followed by a change to 100 % A in 1 min, and an equilibration step for 6 min at 100 % A. For MS analysis, a Thermo QExactive Plus Orbitrap equipped with a HESI ion source running in positive mode was used. For full scan, the resolution was 17.000 with an AGC target of 1e6 and max. IT of 150 ms.

### Zymogram gels

Peptidoglycan isolation from *S. aureus* and *B. subtilis* for zymogram gels was conducted as described before (Fukushima & Sekiguchi, 2016). Briefly, bacterial cultures were grown in LB medium overnight (37°C, 180 rpm), cells were collected by centrifugation (6000 x g, 15 min), washed twice in PBS, and ruptured with glass beads using the Precellysis24 homogenizer (Bertin Technologies). Beads were removed by filtering and cell wall fraction was pelleted by centrifugation (20000 x g, 30 min). The pellet was boiled in 8 % SDS for 10 h, washed three times with PBS, resuspended in PBS, and stored at 4°C. Lytic activity of the CwhA protein was detected using the protocol of de Jonge et al (de Jonge *et al*, 1991). Briefly, proteins were resolved using 12.5 % SDS polyacrylamide gels containing 10 µl/ml isolated cell wall suspension (*S. aureus, B. subtilis*) or 0.2 % lyophilized cells (*M. luteus*). After gel electrophoresis, SDS was removed by washing the gels three times in 50 ml 25 mM Tris-HCl (pH 7) containing 1 % of Triton X-100 within a period of 18 h at room temperature. Gels were stained in 20 ml of staining solution (1 % methylene blue in 0.01 % KOH) for 30 s and immediately destained in H_2_O. The water in the washing bath was changed until a lytic band was clearly visible.

### Live cell imaging of GFP-expressing *S. aureus*

The GFP-expressing *S. aureus* strain (Balwit *et al*., 1994; Kahl *et al*., 2000) was harvested from LB agar plates, a homogenous suspension was prepared in 50 mM Tris buffer (pH 8) and diluted to the optical density of 1.0 at 600nm. 100 µl of the suspension was transferred into single wells of a clear flat-bottom 96-well microplate. After the addition of 5 µl PBS, BSA (bovine serum albumin, 2 mg/ml), or CwhA (2 mg/ml) to selected wells, the plate was transferred to an automated live cell imaging microscope (Celldiscoverer7, Zeiss) and the plate was incubated at 37 °C and 5 % CO_2_. Images were taken every 10 minutes over a time course of 5 hours and the average area fluorescence was determined by Image J for each time point.

### Co-incubation experiments

Fungal mycelia of *A. fumigatus* were harvested from 100 ml liquid AMM cultures that were inoculated with 1.5 x 10^8^ conidia and grown for two days at 37° and 200 rpm. Mycelia were collected using Miracloth (Merck Millipore) and washed with 20 ml sterile ddH_2_0. A small amount of mycelia (∼250 mg) was transferred to a new 500 ml flask containing 100 ml DMEM + 10 % FCS using an inoculation loop. 3 ml of a bacterial suspension in PBS at OD_600_ of 7 were added after growing bacteria overnight in LB medium and washing twice with PBS. The co-cultures were incubated for 4 hours at 37° and 200 rpm, and mycelia were collected using Miracloth (Merck Millipore). The fungal mycelia were washed with 250 ml ddH_2_0, dried using paper towels, flash frozen in liquid nitrogen and stored at -80°C for subsequent RNA isolation and qRT-PCR analysis.

### CwhA-induced bacterial lysis

Bacterial overnight cultures in LB medium were diluted 1:100 in 10 ml fresh LB media and incubated for 3 hours. Cells were then centrifuged at 10,000 g for 5 min and washed two times in PBS. The pellet was resuspended in 50 mM Tris buffer (pH 8) to reach an OD_600_ of 1.0 and 200 µl were transferred to 1.5 ml reaction tubes for CFU counts or to 96-well microplates for automated measurement of the absorbance at 600nm using a microplate reader (TECAN infinites series). 10 µl of CwhA diluted from a 4 mg /ml solution in PBS to final concentrations of 200 - 0.4 µg/ml were added and the samples were incubated at 37°C and 300 rpm for up to 3 hours for CFU counts or without shaking at 37°C for automated reading of OD. For CFU counts, 10 µl of the samples were collected every 15-30 minutes and serial dilutions were plated on LB agar plates. PBS was used as a control.

### Infection of A549 cells for gene expression studies

A549 cells, a human type II alveolar epithelial cell line provided by American Type Culture Collection (ATCC: CCL-185), were seeded in 6-well microplates at 9 x 10^4^ cells/well in DMEM + 10 % FCS (Gibco) and incubated for 24 h at 37 °C, 5 % CO_2_ and 21 % O_2_. Cells were washed once with PBS and 5 ml of a spore suspension containing 9 x 10^5^ conidia in DMEM + 10 % FCS were added to the cells. The plates were incubated at 37°C, 5 % CO_2_ and 21 % O_2_ for 9 h. For growth under hypoxia, plates were transferred to a hypoxystation (H35, Don Whitley Scientific) and incubated for additional 6-24 h at 37 °C, 5 % CO_2_ and 0.2 % O_2;_ for normoxia, plates remained at 21 % O_2_. For the 24 h incubation, the culture medium was replaced with fresh DMEM + 10 % FCS to avoid starvation of cells and remove overgrowing fungi. For RNA extraction, plates were transferred on ice, cells washed with PBS, and host cells lysed with ice-cold ddH_2_O. Attached cells and fungi were removed from the bottom surface using a cell scraper and transferred into 1.5 ml reaction tubes. Fungal mycelia were washed once in ddH_2_O, collected by centrifugation at 4°C, 13,000 g for 5 min, and the pellet was resuspended in 250 µl RL buffer (Roboklon) containing 10 µl/ml mercaptoethanol. The samples were flash-frozen in liquid nitrogen and stored at -80°C until RNA isolation.

### Damage assays

Vegetative cells of *D. discoideum* AX4 were grown in petri dishes containing HL5 media with 1% [w/v] glucose (Formedium) at 22 °C until reaching 80% confluency. Cells were harvested by scraping and counted in the CASY cell counter (OLS Bio) and adjusted to 10^6^ cells/ml in 24 well plates with HL5 media with 1% glucose. The purified protein was added up to a final concentration of 130 µg/ml and the plate was incubated at 22 °C. Live cell numbers were determined with the CASY counter after 0, 1.5, 4, and 26 h of incubation.

A549 cells or MH-S cells (ATCC; CRL-2019™) were seeded in 96 well microplates at 2x 10^4^ cells/well in DMEM (Gibco) or RPMI (Gibco) supplemented with 10 % FCS in a total volume of 200 µl/well. Cells were incubated for 2 days at 37°C, 5 % CO_2_ and 21 % O_2_ to obtain a confluent layer and the medium was exchanged for DMEM for A549 and RPMI for MH-S cells, both containing 1 % penicillin /streptomycin solution (Gibco). Recombinant CwhA protein was added in a volume of 5 µl/well with final concentrations of 25, 12.5, 6.3, and 3.1 mg/ml, and the plates were incubated at 37°C, 5 % CO_2_ and 21 % O_2_ for 24 h. 5 µl of 5 % Triton-X were added to the positive control well and incubated for another 5 minutes. The plate was centrifuged for 10 min at 250 g and 10 µl of the supernatant were transferred to a new microplate containing 90 µl PBS. The reaction mixture of the Roche Applied Science Cytotoxicity Detection Kit was prepared according to the manufacturer’s protocol, and 100 µl of the mix was added to each well and incubated for 30 min at room temperature in the dark.

The absorbance was measured at 492 and 690 nm using a microplate reader (TECAN Infinite series) to detect released lactate dehydrogenase (LDH) as a read-out for cytotoxicity.

### RNA extraction and quantitative RT-PCR

Fungal mycelium was disrupted by glass beads (0.5 mm, Biospec. Products) and RNA was isolated using the GeneMatrix Universal RNA Purification Kit (Roboklon, Berlin) according to the manufacturer’s instructions specified for yeast RNA isolation. RNA was eluted in 30 µl RNase-free water, the concentration was determined with Nanodrop ND-1000 (Thermo Fisher), and aliquots were prepared and stored at -80°C until usage. RNA quality was tested using the RNA 600 Nano Kit and the Bioanalyzer 2100 (Agilent Technologies) according to the manufacturer’s instructions. cDNA was generated from 800 ng of isolated RNA using Superscript III Reverse Transcriptase (Invitrogen). 2xGo Taq qPCR Mastermix (Promega) was used for PCR. Relative gene expression levels were determined by the 2^-ΔΔCt^ method (Schmittgen & Livak, 2008) with *cox5* and *act1* as housekeeping genes. RNA extracted from *A. fumigatus* mycelia grown in DMEM solution served as reference.

### Determination fungal survival in macrophages

Adherent human monocyte-derived macrophages were kindly provided by Marcel Sprenger (Leibniz Institute for Natural Product Research and Infection Biology, Jena, Germany) (Sprenger *et al*, 2020). For infection 5x 10^5^ macrophages/well were seeded in 24 well plates in RPMI supplemented with 10 % FCS in a total volume of 500 µl/well. Cells were allowed to adhere over night at 37°C and 5 % CO_2_ and infected with conidia of *A. fumigatus* wild type and DcwhA at an MOI of 1 in technical triplicates for 12 and 24 h 37°C and 5 % CO_2_. Cells were washed in PBS twice and macrophages were lysed with 500 µl ddH_2_0. Serial dilutions were prepared with PBS and plated onto malt agar plates. CFUs were determined after 2 days of incubation at 37°C. For statistical analysis, the mean of the technical triplicates was used.

### PBMC isolation and stimulation

PBMCs were isolated from human blood that was drawn from healthy donors into 10 ml EDTA tubes. Blood was diluted in PBS (1:1) and cell fractions were separated in lymphocyte separation medium (density 1.077 g/ml, Capricorn Scientific) by density gradient centrifugation according to the protocol supplied by the manufacturer. Cells were washed twice with PBS and resuspended in RPMI 1640^+^. Cells were supplemented with 10 % human serum, plated in 96 well flat bottom plates (Corning, NY, USA) at a final concentration of 2.5 x 10^6^ cells/ml and in a total volume of 200 µl. The stimulation was performed by adding 1 x 10^7^/ml heat-killed *A. fumigatus* conidia, 10 µg/ml Muramyl-dipeptide (MDP, kindly provided by Mark Gresnigt), a combination of heat-killed conidia and MDP, or 25 µl of digest supernatants. To prepare digest supernatants, the following combinations were used: (i) 50 µl peptidoglycan derived from *S. aureus* (*S. aureus* PG), (ii) 50 µl *S. aureus* PG + 20 µg CwhA, (iii) 20 µg CwhA, were incubated for 16h at 37 °C. A combination of MDP and conidia was used as positive (high) control, MDP only was used as a low control. The reaction was stopped by boiling at 98°C for 5 min and non-soluble *S. aureus* PG was removed by centrifugation and transfer of the supernatants to fresh tubes. After cell stimulation for 24 h culture, supernatants were collected and stored at -20°C until cytokine measurement. Experiments were conducted with technical duplicates; for statistical analysis, the mean of the duplicates was used.

### Cytokine measurements

The cytokine levels in stimulated PBMC supernatants were quantified using the uncoated ELISA kits for IL-1β and TNF-α cytokines (GE Healthcare) according to the protocols supplied by the manufacturer.

### Statistical analysis

Statistical analyses were performed with GraphPad version 9.12. Details on the test used for each data set and the number of replicates are stated in the respective figure/table legends.

## Data availability

The authors declare that the data supporting the findings of this study are available within the paper and its Supplementary Information Files.

## Acknowledgements

We thank Maria Stroe, Carmen Schult, and Dirk Femerling for excellent technical assistance, Ralf Ehricht and Stefan Monecke for providing MRSA strains, Sven Krappmann for the plasmid pSK379 and advice on the overexpression strategy, and Mark Gresnigt for encouraging discussions. The work was financially supported by the BMBF (Forschungscampus InfectoGnostics, project IDES, FKZ: 13GW0096E, to IDJ) and the German Research Foundation (DFG; TRR 124 FungiNet, “Pathogenic fungi and their human host: Networks of Interaction,” DFG project number 210879364, Project Z2 to FvE).

## Author contributions

Conception and design of the study was performed by S.M., U.B. and I.D.J. All authors contributed with data acquisition and analysis: S.M. generated the mutants, expressed and purified the protein, isolated bacterial peptidoglycan, tested fungal survival in macrophages and performed cleavage assays, zymogram gels, damage assays, bacterial survival experiments, PBMC isolation and stimulation experiments with subsequent cytokine measurement. U.B. carried out LC-MS analysis of cleavage products. F.Ho. and F.V.E performed MALDI-IMS. Z.F. extracted RNA and fungal protein, and performed fungal survival assays, qRT-PCR, SDS-PAGE and western blots. F.K. performed cell infection, hypoxia and co-incubation experiments, followed by RNA extraction and qRT-PCR. M.F. contributed to fungal cloning and co-incubation experiments. A.N.J.I. and F.Hi. performed amoeba experiments. Data were interpreted by S.M., U.B., A.N.J.I., F.Ho, Z.F., F.Hi., F.K., A.A.B. and I.D.J. S.M., U.B. and I.D.J. wrote and all authors commented on and edited the manuscript.

