## Supplementary Figures, Tables, and information for "Identification of a fungal antibacterial endopeptidase that modulates immune responses"

### Supplementary Figure 1

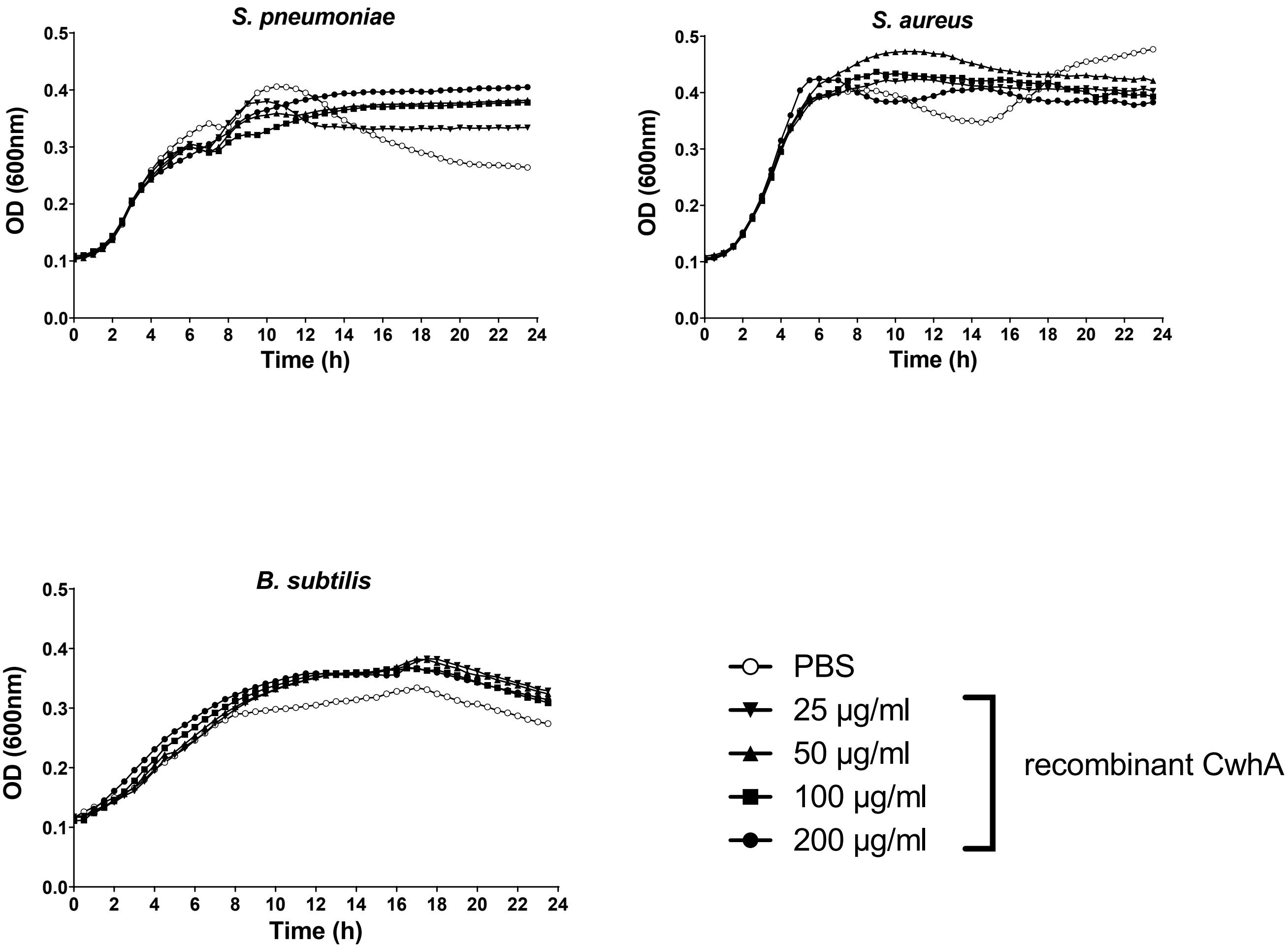


**Supplementary Figure 1:** Bacterial growth in LB media supplemented with CwhA. The optical density at 600nm was measured every 30 min in a plate reader after addition of PBS or different concentrations of recombinant CwhA (25- 200 μg/ml). One representative experiment is shown.

### Supplementary Figure 2

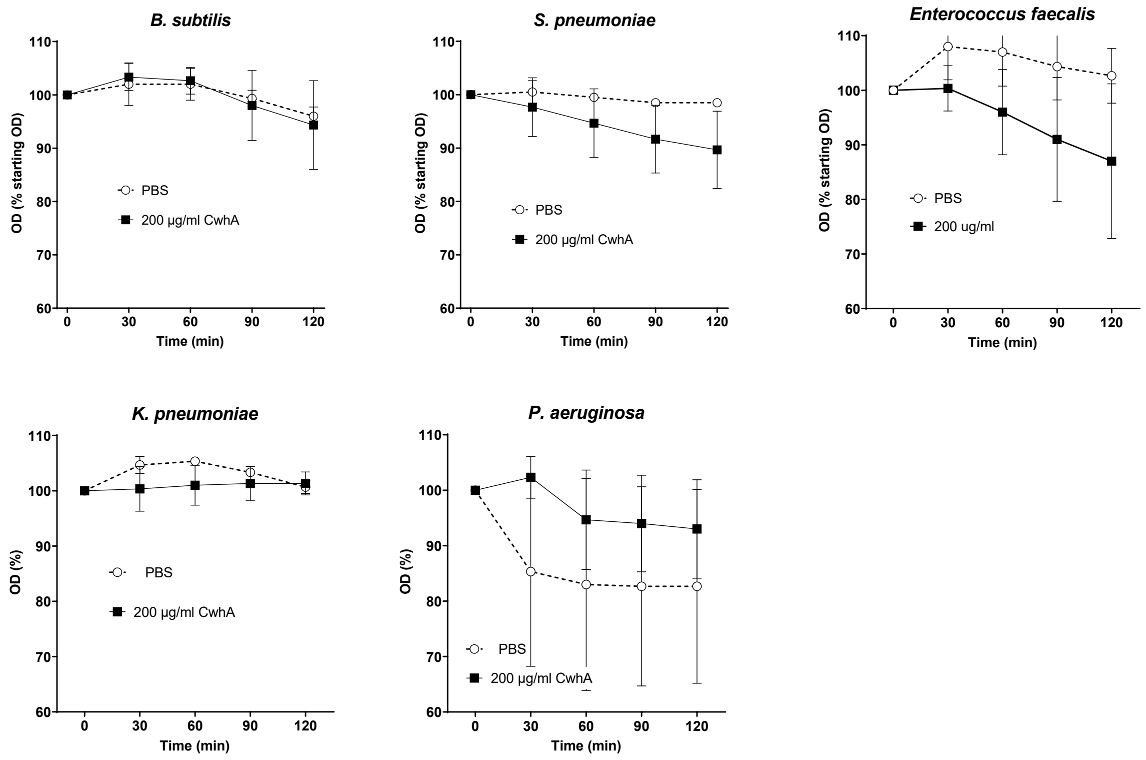


**Supplementary Figure 2. CwhA-mediated lysis of bacterial species.** The optical density (600 nm) after addition of recombinant CwhA (200 µg/ml) to bacterial suspensions (*E. faecalis*, *S. pneumoniae*, *B. subtilis*, *K. pneumoniae* and *P. aeruginosa*) in 50 mM Tris buffer (pH 8) was measured at the indicated time points; three independent experiments, data shown as mean ± SDs. Statistic analysis was performed with 2way ANOVA; treatment had no significant effect.

### Supplementary Figure 3

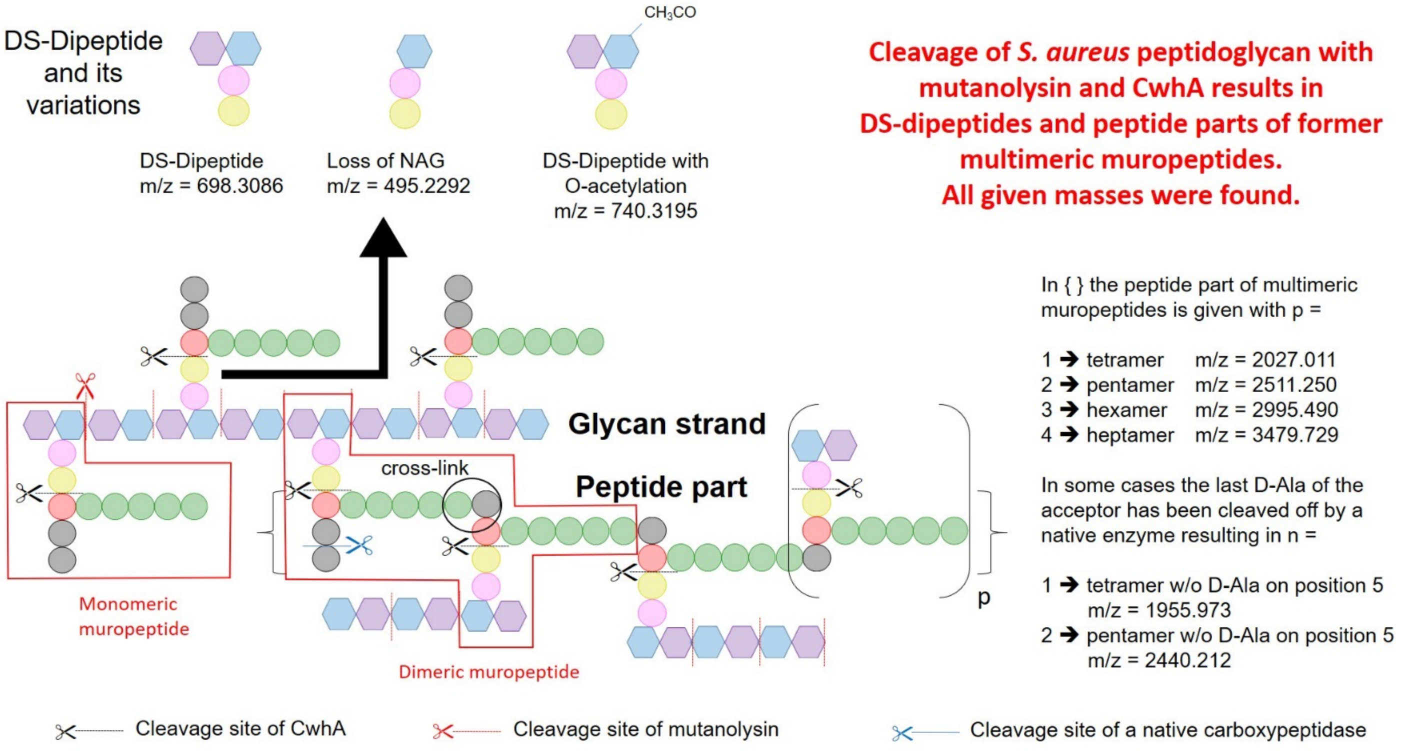


**Supplementary Figure 3. Schematic depiction of peptidoglycan cleavage by mutanolysin and CwhA.**

Supplementary Figure 4

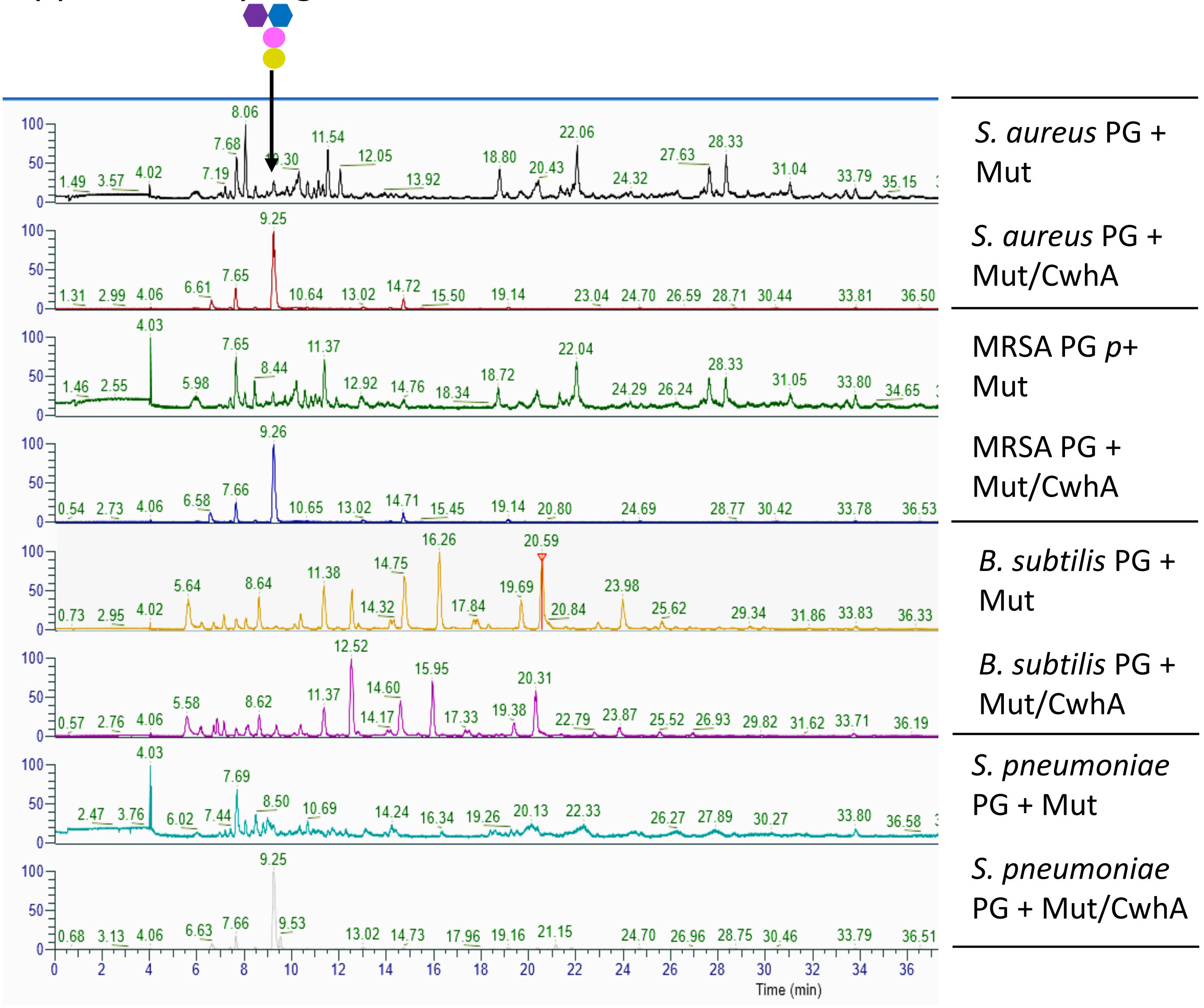


**Supplementary figure 4. Muropeptide profile of peptidoglycan from *S. aureus* SA113, MRSA 181*, S. pneumoniae* (Klein 1884), *B. subtilis* (Ehrenberg 1835) obtained by UPLC/MS after treatment with mutanolysin or/and CwhA.** Masses of peaks are shown in Supplementary table 3. The asterix indicates peaks that are variants of Tetra-Tri.

Supplementary Figure 5

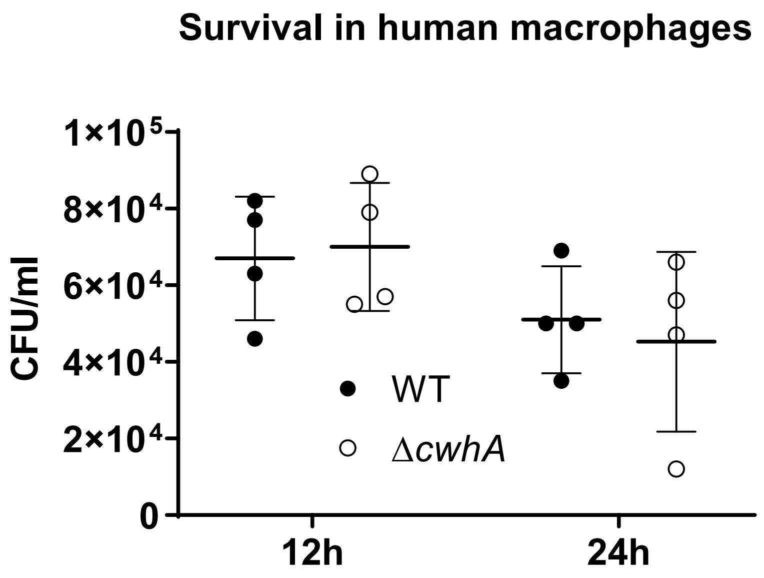


**Supplementary figure 5: Survival of *Aspergillus fumigatus* wt and Δ*cwhA* in human macrophages.**
Differentiated adherent human monocyte-derived macrophages were infected with fungal conidia at an MOI of 1 for 12 h and 24 h and lysed to determine fungal growth on malt agar plates. Data from four independent donors is presented as scatter dot plot with mean ± SD. Analysis by one-way ANOVA and Tukey's multiple comparisons test showed no significant differences between the strains.

**Supplementary Tables**

**Supplementary Table 1**

| **m/z** | **Modification** | **Missed Cleavage** | **Peptide Sequence** | **MSI m/z values with adducts** |
|---|---|---|---|---|
| 1092.378 | none | 1 | (K)ASRIPPLPLK(L) | 1093.38558 [M+H] |
| 1177.397 | none | 1 | (R)TVFTPGIKASR(I) | 1178.40468 [M+H] |
| 2017.331 | none | 2 | (R)HVPYSMNKVYPDPQGRK(Y) | 2018.33858 [M+H] |
| 1156.301 | none | 1 | (K)KEYNTLECR(G) | 1174.34443 [M+NH4] |
| 1150.437 | none | 1 | (-)MRTVFTPGIK(A) | 1173.42652 [M+Na] |
| 1061.277 | 1Met-loss+Acetyl | 1 | (-)MRTVFTPGIK(A) | 1125.29277 [M+ACN+Na] |

**Supplementary Table 1:** Peptides predicted by *in silico* digestion and identified by MALDI-imaging mass spectrometry (MSI) in both lung sections and control spots containing recombinant CwhA. Distribution of these peptides closely matched the example of peptide (K)KEYNTLECR(G) shown in Figure 1.

**Supplementary Table 2**

| Strain | PBS | CwhA | *P* value |
|---|---|---|---|
| ATCC13420 | 92.9 ± 1.7 | 74.7 ± 1.1 | 0.0078 |
| MRSA 124 722 | 92.0 ± 0.0 | 68.0 ± 1.7 | 0.0017 |
| MRSA 124 622 | 93.7 ± 5.8 | 62.0 ± 1.7 | 0.0054 |
| MRSA124 737 | 93.7 ± 0.0 | 70.0 ± 3.5 | 0.0128 |
| MRSA 293 888 | 93.0 ± 1.7 | 68.7 ± 4.0 | 0.0075 |
| MRSA 94 003 | 96.0 ± 1.7 | 78.7 ± 1.1 | 0.0004 |
| MRSA 314 432 | 91.0 ± 2.6 | 77.7 ± 4.7 | 0.0857 |
| MRSA 124 940 | 90.7 ± 7.5 | 71.3 ± 9.2 | 0.1835 |
| MRSA 124 411 | 92.0 ± 1.7 | 72.7 ± 11.5 | 0.1278 |
| MRSA 124 328 | 92.7 ± 4.0 | 69.7 ± 11.0 | 0.0289 |

**Supplementary Table 2:** Relative optical density (600 nm, normalized to 0 min) of *S. aureus* strains after 120 min incubation in PBS or 100 mg/ml CwhA. Mean ± SD of values obtained in three independent experiments; p value: Comparison of PBS and CwhA treatment by two-tailed paired t test. *P* values < 0.05 are highlighted by a light grey background.

**Supplementary Table 3**

| Peak | Measured $[M+H]^+$ | Basic structure | Variations | | | |
|---|---|---|---|---|---|---|
| | | | GlcNac missing | additional O-Acetylation | amidated mDpm | N-Acetylation missing |
| *S. aureus* | | | | | | |
| 1 | 968.4781 | DS-Pentapeptide | | | | |
| 2 | 1253.5867 | 1xDS-Pentapeptide-Gly5 | | | | |
| 3 | 2417.1157 | 2xDS-Pentapeptide-Gly5 | | | | |
| 4 | 3580.6537 | 3xDS-Pentapeptide-Gly5 | | | | |
| 5 | 2328.0688 | Cyclic 2xDS-Tetrapeptide-Gly5 | | | | |
| P | 2027.0106 | Peptide part of a tetramer | | | | |
| P | 2511.2478 | Peptide part of a tetramer | | | | |
| P | 2995.4873 | Peptide part of a tetramer | | | | |
| P | 3479.72915 | Peptide part of a tetramer | | | | |
| Dx | 495.2292 | 1xDS-Dipeptide | 1 | | | |
| D1 | 698.3086 | 1xDS-Dipeptide | | | | |
| Dy | 740.3195 | 1xDS-Dipeptide | | 1 | | |
| D2 | 1375.58432 | 2xDS-Dipeptide | | | | |
| D3 | 2052.85794 | 3xDS-Dipeptide | | | | |
| D4 | 2730.13284 | 4xDS-Dipeptide | | | | |
| D5 | 3407.41374 | 5xDS-Dipeptide | | | | |
| D6 | 4084.68696 | 6xDS-Dipeptide | | | | |
| D7 | 4761.97016 | 7xDS-Dipeptide | | | | |
| D8 | 5439.25176 | 8xDS-Dipeptide | | | | |
| D9 | 6116.53188 | 9xDS-Dipeptide | | | | |
| *B. subtilis* | | | | | | |
| 1 | 870.3922 | DS-Tripeptide | | | 1 | |
| 2 | 871.3762 | DS-Tripeptide | | | | |
| 3 | 1750.7930 | DS-Tetrapeptide - DS-Tripeptide | | | 2 | 1 |
| 4 | 1751.7770 | DS-Tetrapeptide - DS-Tripeptide | | | 1 | 1 |
| 5 | 1792.8042 | DS-Tetrapeptide - DS-Tripeptide | | | 2 | |
| 6 | 1752.7612 | DS-Tetrapeptide - DS-Tripeptide | | | | 1 |
| 7 | 1793.7890 | DS-Tetrapeptide - DS-Tripeptide | | | 1 | |
| 8 | 1793.7878 | DS-Tetrapeptide - DS-Tripeptide | | | 1 | |
| 9 | 1794.7544 | DS-Tetrapeptide - DS-Tripeptide | | | | |
| Dz | 657.2823 | DS-Dipeptide | | | | 1 |
| D1 | 699.2931 | DS-Dipeptide | | | | |

**Supplementary Table 3:** Masses of peptides shown in Figure 4.

**Supplementary Table 4**

| Strain no. | Strain type | Origin |
|---|---|---|
| 124 722 | CC5/ST228-MRSA-I, Süddeutscher | human |
| 124 622 | CC133-MSSA (lukF-P83/lukM+) | Veterinär |
| 124 737 | CC8-MRSA-IVh/j (sea+), "UK-EMRSA-2" | human |
| 293 888 | CC398-MRSA-[V/VT+ccrB1] | veterinary |
| 94 003 | CC5-MRSA-II (tst1+), New York-Japan Clone | Mu50 / reference strain |
| 314 432 | CC772-MRSA-V/VT (PVL+) | human |
| 124 940 | CC80-MRSA-IVc (PVL+) | human |
| 124 411 | CC8-MRSA-[IV+ACME] (PVL+), USA300 | human |
| 124 328 | CC22-MRSA-IV, UK-EMRSA-15/Barnim EMRSA | human |

**Supplementary Table 4**: Multiresistant *S. aureus* (MRSA) strains used in this study. Strains are part of the strain collection of the University Hospital Dresden, Germany, and were kindly provided by S. Monecke.

**Supplementary Table 5**

| Name | Sequence 5' → 3' |
|---|---|
| Del_p60A_F1 | CACGACGTTGTAAAACGACGGCCAGTGCCAGGTATAGCGAGGTTGCATCC |
| Del_p60A_R2 | GAGGCCATCTAGGCCATCAAGCGCGAGCAGAGTTGACAAGTG |
| Del_p60A_F3 | GGCCTGAGTGGCCATCGAATTCGATGGAGCGTTGGATAGGTG |
| Del_p60A_R4 | GATCCTCTAGAGTCGACCTGCAGGCATGCAGAAGCCATTCTCTGCTATTC |
| ptrA_for_II | GAATTCGATGGCCACTCAGGCC |
| ptrA_rev_II | GCTTGATGGCCTAGATGGCCTC |
| o_p60AFor | GAGGATCCGTACCCCATCACTGGCAACG |
| o_p60ARev | GAGAATTCTTAGTCCACAACGCG |
| Expr_p60A_for | ATGCGCACTGTATTCACTCC |
| Expr_p60A_rev | CATGATATCTTAGTCCACAACGCGGATGTA |
| RT_p60Afor | TGGCTTCAGCACTGTCACTC |
| RT_p60Arev | TGTACTTGACGCAGCCGTAG |
| RT_Cox5 for | ATCTGTTCGCCAAGCCCAAG |
| RT_Cox5 rev | TCACTGCTGACACCGTAGAG |
| RT_act1_for | CCACGTCACCACTTTCAACTC |
| RT_act1_rev | CTGCATACGGTCGGAGATAC |

**Supplementary Table 5**: Oligonuceotides used in this study.

#### General features of peptidoglycan

Most eubacteria are surrounded by the so-called peptidoglycan (PG) sacculus that is responsible for cell shape and counteracts the internal pressure [1]. It is a macromolecule composed of glycan strands that are cross-linked by short peptides. Alternating N-acetylglucosamine (NAG) and N-acetylmuramic acid (NAM) are β-1,4 linked to form glycan strands of variable length. Attached to the lactyl group of MurNAc is a stem peptide of five amino acids with a quite conserved sequence: L-alanine – D-iso-glutamate – L-lysine – D-alanine – D-alanine for Gram-positive organisms and meso-Diaminopimelic acid (mDpm) on position three of Gram-negatives and also of *Bacillus* species [2]. There are several possible modifications known. For example, glycan strands can be O-acetylated, O-deacetylated or N-deacetylated. These modifications often result in lysozyme resistance [3-7]. Cross-linking of the glycan strands occurs between two adjacent stem peptides by formation of a peptide bond between D-alanine on position four and the amino acid on position three (L-Lys or m-Dpm) catalyzed by the penicillin-binding proteins and release of the fifth amino acid D-alanine [8,9]. Cross-linking can be either direct or indirect by an interpeptide bridge, *i.e.* the penta-glycine bridge in *S. aureus* [2]. For analysis by HPLC PG is isolated and digested by the muraminidase mutanolysin into muropeptides thereby cleaving the glycosidic bond [10-13]. The resulting disaccharide units can still be cross-linked thereby forming dimers, trimers and multimers whose retention times increase [14]. Digestion of the PG macromolecule into smaller units also occurs in nature and is a defense mechanism of many organisms. Resulting small muropeptides that miss the NAG-part are immunogenic. The Muramyl-dipeptide is sensed by NOD2, the m-Dpm containing Muramyl-tripeptide (MTP) is recognized by NOD1 [15-17].
